# The Bardet-Biedl Syndrome complex component BBS1 regulates proteasome-dependent F-actin clearance from the centrosome to enable its translocation to the T cell immune synapse

**DOI:** 10.1101/2020.11.30.403980

**Authors:** Chiara Cassioli, Anna Onnis, Francesca Finetti, Nagaja Capitani, Ewoud B Compeer, Micheal L Dustin, Cosima T Baldari

## Abstract

Components of the intraflagellar transport (IFT) system that regulates the assembly of the primary cilium are exploited by the non-ciliated T cell to orchestrate polarized endosome recycling to sustain signaling during immune synapse formation. Here we have investigated the potential role of BBS1, an essential core component of the Bardet-Biedl syndrome complex that cooperates with the IFT system in ciliary protein trafficking, in the assembly of the T cell synapse. We show that BBS1 allows for centrosome polarization towards the immune synapse by promoting its untethering from the nuclear envelope. This function is achieved through the clearance of centrosomal F-actin and its positive regulator WASH, a process that we demonstrate to be dependent on the proteasome. We show that BBS1 regulates this process by coupling the 19S proteasome regulatory subunit to the microtubule motor dynein for its transport to the centrosome. Our data identify the ciliopathy-related protein BBS1 as a new player in T cell synapse assembly that acts upstream of the IFT system to set the stage for polarized vesicular trafficking and sustained signaling.

## INTRODUCTION

Vesicular trafficking has emerged over recent years as a central factor in the assembly and function of the immune synapse (IS), a specialized membrane domain that T cells forms at the interface with antigen presenting cells (APC) carrying specific MHC-bound peptide antigen [1]. Essential components of the T cell receptor (TCR) signaling cascade are present as two pools with different and dynamic subcellular localization These include the TCR itself, the initiating kinase Lck and the transmembrane adaptor LAT that couples TCR engagement to multiple intracellular signaling modules [2–4]. One pool is associated to the plasma membrane and is readily mobilized upon TCR triggering to orchestrate the signals that initiate the topological reorganization of receptors and signaling mediators leading to the generation of the mature IS architecture. The second pool is associated to recycling endosomes, which undergo polarized exocytosis at the IS following centrosome translocation towards the T cell:APC interface to sustain signaling [5–7]. Vesicular trafficking also regulates the fate of exhausted TCRs, which accumulate at the IS center and undergo endocytosis to be sorted for recycling or lysosome-mediated degradation [8–10]. Post-endocytic TCRs are also exploited for intercellular communication through their incorporation into synaptic ectosomes that are released in the synaptic cleft to be taken up by the cognate APC [11].

While endosome recycling had long been known for the TCR and, based on its long half-life, viewed as a mechanism of quality control [10], accumulating evidence for a crucial role of membrane trafficking in IS assembly and function has placed the focus on the characterization of the underlying molecular machinery. We identified as an unexpected new player in synaptic trafficking the intraflagellar transport (IFT) system. This multimolecular complex, consisting of the IFT-A and IFT-B subcomplexes, regulates the growth, disassembly and signaling function of the primary cilium, a small appendage present on the majority of mammalian cells, a major exception being hematopoietic cells [12]. In particular, we found that IFT20 promotes IS assembly together with other IFT-B components by regulating polarized TCR recycling in the non-ciliated T cell [13,14]. Many other ciliary proteins have since been demonstrated to be co-opted by T cells for IS formation, including adaptors (e.g. Unc-119, the Rac1 interacting protein FIP3, the microtubule-binding protein EB-1), small GTPases (e.g. Rab8, Rab29, Arl3, Arl13) and tethering proteins (e.g the SNAREs VAMP-3 and VAMP-7, the golgin GMAP210) [3,15–21]. Additionally, ciliary signaling pathways, such as the Hedgehog signaling cascade and lipid kinase and phosphatase networks, are exploited by cytotoxic T cells for the assembly of the lytic synapse [22,23], leading to the hypothesis that the IS and the primary cilium are functional homologues [24–26].

Here we have investigated the potential role of the Bardet-Biedl syndrome complex or BBSome, a key regulator of ciliary function that functionally and physically interacts with IFT-B, in IS assembly in the non-ciliated T cell. The BBSome is an octameric, centrosome-associated complex that undergoes IFT, retrieving membrane cargoes such as activated GPCRs at the tip of the cilium, where it assembles with IFT-B to promote their exit from the cilium by regulating their passage through the diffusion barriers at the ciliary base [27,28]. We show that the BBSome core component BBS1 [29] is essential for centrosome translocation towards the T cell interface with cognate APC. We provide evidence that BBS1 promotes the proteasome-dependent F-actin clearance from the centrosome, on which its detachment from the nuclear membrane and mobilization to the IS crucially depend. We show that BBS1 regulates this process by coupling the 19S regulatory subunit of the proteasome to the microtubule motor dynein for its transport to the centrosome. Our data identify the ciliopathy-related protein BBS1 as a new player in the assembly of the T cell IS that acts upstream of the IFT system to set the stage for polarized recycling and sustained signaling.

## MATERIALS AND METHODS

### Cells, plasmids and antibodies

Cells included Jurkat T cells, Raji B cells and BJ-5at fibroblasts. Primary T cells were isolated from peripheral blood of healthy donors by negative selection using RosetteSept^™^ Human T Cell Enrichment Cocktail (STEMCELL Technologies, 15061) and activated with Dynabeads^®^ Human T-Activator CD3/CD28 (ThermoFisher Scientific, #111.13D) for 48 hours. B and T lymphocyte cell lines as well as primary T cells were grown at 37°C, 5% CO_2_, in RPMI-1640 Medium (Merck, #R8758) supplemented with 10% iron-enriched bovine calf serum (BCS; GE Healthcare HyClone, SH30072.03) and 50 U/ml of human IL-2 IS (Miltenyi Biotec, #130-097-142) for primary T cells. BJ-5at fibroblasts were kept in a 4:1 mixture of the Dulbecco’s Medium (Merck, #D6429) and Medium 199 (Gibco, #31153) supplemented with 10% fetal bovine serum (FBS; Euroclone, #ECS0180L) and 0.01 mg/ml hygromycin B (Invitrogen, #10687010).

Plasmid included Mission^®^ pLKO-puro Non-Target shRNA (Sigma-Aldrich, #SHC016V), pLKO shRNA targeting BBS1 (Sigma-Aldrich, #TRCN0000417688), pEGFP-N1-hBBS1. To construct a C-terminal GFP-tagged BBS1 (BBS1-GFP), CDS sequence of BBS1 was amplified from pCS2-Myc6-BBS1 construct [28] using the primers described in Table S1 and the PCR product subcloned into pEGFP-N1 [30]. All primary commercial antibodies used in this work are listed in Table S2, together with information about the dilutions used for immunoblotting, immunofluorescence, immunoprecipitation and flow cytometry. Secondary peroxidase-labeled antibodies were from Jackson ImmunoResearch Laboratories (anti-mouse, #115-035-146; anti-rabbit, #111-035-14), and Alexa Fluor 488-, 555- and 647-labeled secondary antibodies were from ThermoFisher Scientific (anti-mouse 488, #A11001; anti-rabbit 488, #A11008; anti mouse 555, #A211422; anti-rabbit 555, #A21428; anti-mouse 647, #A21236).

### Stable BBS1-depleted Jurkat T cell line and transient transfections

A stable BBS1-depleted Jurkat T cell line was generated by transducing cells with lentiviral particles carrying a non-targeting control shRNA (ctr) or a shRNA specific for BBS1 (J KD) and selecting in puromycin-containing medium at the final concentration of 3.5 μg/ml (Merck, #P8833). Knocked-down Jurkat cells were routinely checked for BBS1 expression by immunoblotting. In order to generate a stable transfectant expressing BBS1-GFP, Jurkat cells were transfected cells by electroporation (Gene Pulser II, Bio-Rad) and monoclonal cell lines were selected in Geneticin Selective Antibiotic (G418 Sulfate; ThermoFisher Scientific, #11811-031)-containing medium at a final concentration of 2 mg/ml. Cell clones expressing BBS1-GFP at the highest levels were screened by flow cytometry and routinely checked by immunoblotting and immunofluorescence. Equal number of cells form the 3 highest expressing clones were pooled for experiments. Transient transfection of Jurkat cells with the BBS1-GFP construct (DNA: cells *ratio* = 1 μg/10^6^ cells) were carried out by using a modification of the DEAE-dextran procedure as described [31].

### Gene-editing of Jurkat cells and activated primary T cells by using CRISPR-Cas9 technology

Specific small guide RNAs (sgRNAs) directing the nuclease Cas9 to BBS1 gene (Table S1) were designed using the web-based tool CRISPOR [32]. For gene-editing of Jurkat T cells, gRNAs were cloned into the pSpCas9(BB)-2A-GFP (PX458) plasmid (a gift from F. Zhang; Addgene, #48138) as described elsewhere [33] and Jurkat cells were transfected with either empty vector or the gRNA-encoding construct using the Nucleofector Solution 2M (5 mM KCl, 15 mM MgCl_2_, 15 mM HEPES, 150 mM Na_2_HPO_4_/NaH_2_PO_4_ pH 7.2, 50 mM Mannitol) [34] and the Amaxa Nucleofector II system (Lonza), Program X-005. Green Fluorescent Protein (GFP)-expressing cells were sorted, subcloned, and screened by immunoblotting. For gene-editing of primary T cells, sgRNA transcription templates were prepared by PCR amplification using the PX458 construct as a template and the primers listed in Table S1, then transcribed *in vitro* using HiScribe™ T7 High Yield RNA Synthesis Kit (NEB, #E2040S). sgRNAs were purified by RNA Clean & Concentration™ (Zymo Research, #R1017). Freshly isolated T cells were activated by incubation with anti-CD3/CD28 Dyneabeads for 48 hours in complete RPMI-1640 medium. Activated cells were transfected using the Human T cell Nucleofector Kit (Amaxa Biosystem, #VPA-1002) and the Amaxa Nucleofector II system (Lonza), Program T-023 with ribonucleoprotein complexes, which were formed by mixing 5 μg of Alt-R^®^ S.p. Cas9 Nuclease V3 protein (IDT, #1081059) and 3 μg of each sgRNA. Cells were allowed to recover in complete RPMI-1640 medium supplemented with 500 U/ml of human IL-2 for 72 hours and then tested for gene-editing by immunoblotting.

### Conjugate formation

In IS experiments, Raji B cells, used as APCs, were loaded with 10 μg/ml of Staphylococcal superantigens A (SEA; Toxin Technologies AT101), B (SEB; Toxin Technologies BT202) and E (SEE; Toxin Technologies ET404) for 2 hours and labelled with 10 μM Cell Tracker Blue for the last 20 min of the incubation with superantigens (SAgs). In particular, SEE was used for Jurkat cells that express TCR Vβ8, whereas a combination of SEA, SEB and SEE were used for primary T cells to cover a substantial proportion of the TCR Vβ repertoire compared with SEE alone. Antigen-independent conjugates of T cells with unloaded Raji B cells were used as negative control.

SAg-loaded or unloaded Raji B cells were washed twice, mixed with Jurkat cells or primary T cells (1:1) and conjugates analyzed at the indicated time points after conjugate formation (i.e. 5 and 15 min). Samples were seeded onto poly-L-lysine (Merck, #P1274)-coated slides (ThermoFisher Scientific, #X2XER208B) and fixed for 10 min in methanol at −20 °C or for 15 min with 4% paraformaldehyde/PBS at room temperature. To image the 19S RP at different time points after conjugate formation (i.e. 0, 1, 2.5 and 15 min) Raji B cells were seeded on poly-L-lysine-coated slides and allowed to adhere for 15 min at 37°C, then T cells were added and slides incubated at 37°C for the indicated time points before fixation.

To assess the role of the proteasome in IS formation, Jurkat or primary T cells were pre-treated for 2 hours with two different proteasome inhibitors, MG132 (10 μM; Sigma-Aldrich, #M7449) or epoxomicin (10 μM; Abcam, #ab144598), prior to conjugate formation. None of the treatments affected cell viability at the concentrations and times chosen for the analyses, as assessed by Trypan blue (Sigma-Aldrich, #T8154) exclusion (Fig. S7B).

### Immunofluorescence acquisition and analysis

Following fixation, samples were washed in PBS for 5 min and stained with primary antibodies overnight at 4°C. After washing with PBS, samples were incubated for 45 min at room temperature with Alexa Fluor 488- and 555-labeled secondary antibodies and mounted with 90% glycerol/PBS.

Confocal microscopy was carried out on a Zeiss LSM700 using a 63x/1.40 oil immersion objective. Images were acquired with pinholes opened to obtain 0.8 μm-tick sections. Detectors were set to detect an optimal signal below the saturation limits. Images were processed with Zen 2009 image software (Carl Zeiss, Jena, Germany). Colocalization analyses were carried out on medial optical sections of either single cells using ImageJ and the JACoP plugin to calculate Manders’ coefficient M_1_, which indicates the proportion of the green signal coincident with a signal in the red channel over its total intensity, and M_2_, which is defined conversely for red [35]. Manders’ coefficients range from 0 to 1, corresponding to non-overlapping images and 100% colocalization between both images, respectively.

In conjugates of T cells with SAg-loaded versus unloaded Raji B cells relative distances of the centrosome from either the contact site with the APC [36] or the nuclear membrane [37] were measured using ImageJ (Fig.S2). Scoring of conjugates for clustering of γ-tub, CD3 and P-Tyr clustering at the IS was based on the presence of the respective staining solely at the T-cell–APC contact site. Recruitment index was calculated for each marker as the *ratio* of CD3 and PTyr fluorescence intensity at the synaptic area, which is manually defined at the T cell:APC contact site, to the entire cell by using ImageJ. Values above 1 indicate an accumulation of the marker at the IS area compared to the entire cell, whereas values below 1 indicate a depletion of the marker at the IS area compared to the entire cell. Colocalization analyses on conjugates were carried out by calculating Manders’ coefficients M_1_ and M_2_ in a 4.5 μm (Jurkat cells) or 2.85 μm (primary T cells) diameter circle centered around the centrosome (Fig.S2).

### RNA purification and RT-PCR

RNA was extracted from Jurkat, primary T cells and BJ-5at fibroblasts using the RNeasy Plus Mini Kit (Qiagen. #74136), reverse transcribed to first-stand cDNAs using iScript™ cDNA Synthesis Kit (Bio-Rad, #1708891) and analyzed by RT-qPCR on 96-well optical PCR plates (Sarstedt) using the SsoFast™ EvaGreen^®^ Supermix (Bio-Rad, #1725204) and specific primers for human transcripts listed in Table S1. HPRT1 was used as a house-keeping gene to normalize transcript levels.

### Cell lysis, immunoblots and co-immunoprecipitation (co-IP) experiments

Cells (2×10^6^/sample for immunoblot analysis on total cell lysates) were lysed in 1% (v/v) Triton X-100 in 20 mM Tris-HCl (pH 8), 150 mM NaCl in the presence of Protease Inhibitor Cocktail Set III (Calbiochem^®^, #539-134) and 0.2 mg Na orthovanadate/ml for 5 min on ice. Protein extracts were quantified with a BCS Total Protein Assay Kit (Euroclone, #EMP014500) and denatured in 4X Bolt LDS Sample Buffer (Life Technoogies, #B0007) supplemented with 10X Bolt™ Sample Reducing Buffer (Life Technologies, #B009) for 5 min at 100°C. Proteins (10 μg) were separated on Bolt Bis-Tris Plus gels (Invitrogen™) or home-made gels and transferred to nitrocellulose or polyvinylidene difluoride membranes (GE HealthCare, #10600023) under wet conditions. Blocking were performed in 5% non-fat dry milk in PBS 0.02% Tween-20. Membranes were incubated in primary antibodies for 1-3 hours at room temperature or overnight at 4°C followed by incubation in 20 ng/ml secondary horseradish-peroxidase (HRP)-conjugated secondary antibodies (Jackson ImmunoResearch Laboratories) for 45 min at room temperature. Secondary antibodies were detected by using SuperSignal™ West Pico Plus Chemiluminescent Substrates (Life Technologies, #34578). Membranes were stripped with ReBlot Plus Mild Antibody Stripping Solution 10X (Chemicon^®^, #2502) and reprobed with a primary antibody. For quantification, blots were scanned using a laser densitometer (Duoscan T2500, Agfa) and densitometric levels were measured using ImageJ software.

For co-IP experiments cells (8-10×10^6^ cells/sample) were lysed in 0.5% (v/v) Triton X-100 in 20 mM Tris-HCl (pH 8), 150 mM NaCl in the presence of Protease Inhibitor Cocktail Set III (Calbiochem^®^, #539-134) and 0.2 mg Na orthovanadate/ml for 5 min on ice. Post-nuclear supernatants were incubated with 3 μg Protein A Sepharose CL-4B (PAS; Ge HealthCare, #GEH17078001) for 1 hour and then with 3 μg PAS plus primary antibody (0.4 ug anti-GFP mAb, 1.5 ug anti-Rpt2, 2 ug anti-γ-tubulin) for 2 hours. PAS controls, which corresponds to the pre-clearing of the lysates with the same amount of PAS used for the co-IP, and PAS-antibody complexes were pelleted and washed 4 times in IP lysis buffer. The immunoprecipitates were eluted in 2X Bolt LDS Sample Buffer supplemented with 10X Bolt™ Sample Reducing Buffer (Life Technologies, #B009), boiled for 5 min and resolved by SDS-PAGE. A fraction (10 μg) of the lysates used for co-IPs were run on the same gel to identify the migration of the specific immunoreactive bands.

### Flow cytometry

Flow cytometry analysis of surface CD3 was carried out on control or BBS1 KD/KO Jurkat cells incubated on ice for 30 min with PE-labelled anti-human CD3ɛ (OKT3; Table S2). Samples were acquired with Guava Easy Cyte cytometer (Millipore) and plotted using FlowJo software (TreeStar Inc., Ashland, OR, USA).

In the reconstitution experiments the percentage of live GFP^+^/propidium iodide^−^ cells was determined by flow cytometry by labelling transfected cells with 0.5 μg/ml of propidium iodide (Sigma-Aldrich, #537059).

Protein tyrosine phosphorylation was analyzed by flow cytometry in conjugates of control or BBS1 KO Jurkat cells and SEE-pulsed Vybrant™ DiO (ThermoFisher Scientific, #V22886)-labelled Raji cells (APC) at different time points after conjugate formation. Conjugates were co-stained with an anti-PTyr antibody (Table S2) and DiO-negative cells were analyzed.

### Statistics and reproducibility

Each experiment was performed ≥ 3 independent times. The exact number of repeats and the number of cells analysed is specified in figure legends. Statistical analyses were performed with Prism software (GraphPad Software). Values with normal distribution were analysed with Student’s *t*-test (paired or unpaired) and one-sample *t*-test (theoretical mean = 1), whereas values without Gaussian distribution were analysed with Kruskal-Wallis test or Mann-Whitney for multiple comparisons. Statistical significance was defined as: ns *P* > 0.05, * *P* ≤ 0.05, ** *P* ≤ 0.01, *** *P* ≤ 0.001, **** *P* ≤ 0.0001.

## RESULTS

### BBS1 localises at the pericentrosomal compartment and accumulates at the T cell IS

To investigate the role of the BBSome in IS assembly in human T cells we first measured the expression of its eight core components (BBS1, BBS2, BBS4, BBS5, BBS7, BBS8, BBS9 and BBS18) in both Jurkat and primary peripheral blood T cells, using as a positive control the ciliated fibroblast cell line BJ-5at. Expression was comparable or higher in T cells compared to ciliated cells, as assessed by RT-qPCR (Fig.S1a). T cell expression was confirmed by immunoblot for BBS1 (Fig.1a), which we chose for further characterization as this protein is essential for recruitment of GTP-bound BBS3/Arl6 into the BBSome to enable its entry into the cilium [29] and has been recently implicated in BBSome stability [38].

**Fig. 1.**
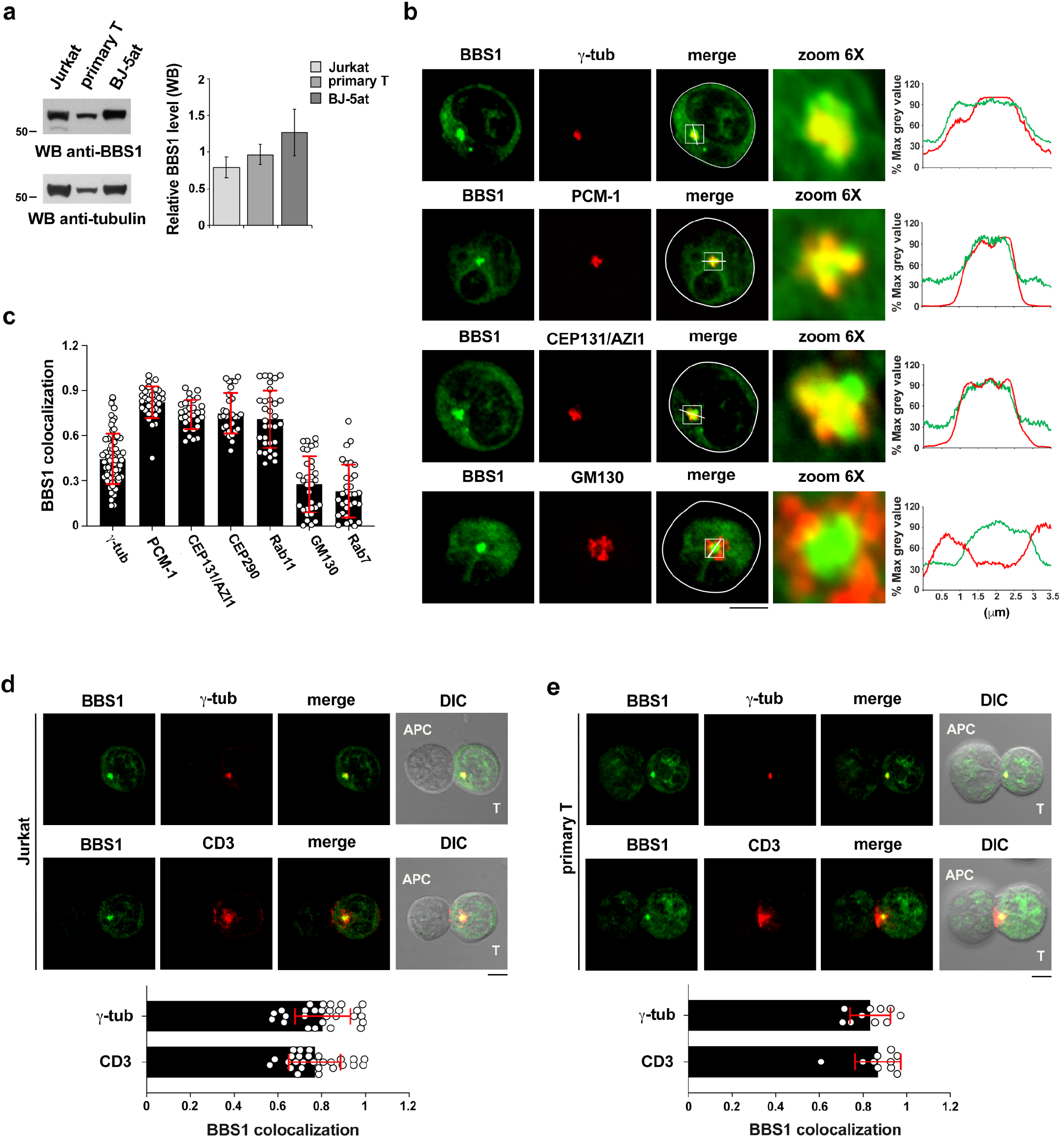
BBS1 localises at the pericentrosomal compartment and accumulates at the T cell IS. (**a**) *Left,* Immunoblot analysis of BBS1 in lysates of Jurkat and primary T cells, using ciliated BJ-5at cells as positive control (n≥3). Tubulin was used as loading control. The immunoblot shown is representative of 3 independent experiments. The migration of molecular mass markers is shown for each filter. The quantification of the immunoblots is shown on the right. (**b**) Immunofluorescence analysis of BBS1-GFP-expressing Jurkat cells costained for γ-tubulin (centrosome), PCM-1 and CEP131/AZI1 (centriolar satellites markers), or GM130 (Golgi apparatus). The histograms show the intensity profiles along the white lines within the selected area in the overlay images for each channel. The raw pixel intensity signals were normalized to maximum intensity pixel of each channel (% max grey value). (**c**) Quantification using Mander’s coefficient of the weighted colocalization of GFP signal with the indicated markers in Jurkat cells transiently or stably transfected with a construct encoding BBS1-GFP. At least 20 cells were analyzed for each marker (n≥3). Representative images (medial optical sections) are shown in (**b**) and (**S2a**). (**d,e**) Immunofluorescence analysis of γ-tubulin and CD3ζ localization in 15-min antigen-specific conjugates of Jurkat cells (**d**) or primary T cells (**e**) transfected with a construct encoding BBS1-GFP and SEE- or SAg-loaded Raji B cells (APC), respectively. A quantification using Mander’s coefficient of the weighted colocalization of the GFP signal with the indicated markers is shown in the bottom panel. The distance of BBS1-GFP (μm) from the T cell:APC contact site, measured from the point of maximal intensity of the GFP signal, was 2.99+/− 0.85 in Jurkat cells (20 cells/sample, n=3) and 1.13+/−0.85 in primary T cells (10 cells/sample, n=2). Size bar, 5 μm. The data are expressed as mean±SD.

The subcellular localization of BBS1 was determined by confocal immunofluorescence analysis of Jurkat cells transfected with GFP-tagged BBS1 (BBS1-GFP) and co-stained with markers of intracellular compartments. Based on the basal body localization of the BBSome in ciliated cells [27,39], these included markers of the centrosome (γ-tubulin) and of the pericentrosomal (PCM-1, CEP131/AZI1, CEP290) and endocytic recycling (Rab11) compartments. Markers of the Golgi apparatus (GM130) and late endosomes (Rab7) were also included. Similar to ciliated cells [40], BBS1 co-localized most prominently with the centrosomal and pericentrosomal compartments, as well as with recycling endosomes (Figs.1b,c,S2a).

During IS formation the centrosome moves towards the APC, allowing for the polarized delivery to the synaptic membrane of a pool of TCR:CD3 complexes associated with recycling endosomes [5,7]. BBS1 was found to polarize to the IS formed in conjugates of Jurkat cells with Raji cells (used as APCs) pulsed with Staphylococcal enterotoxin E (SEE). At the IS BBS1 co-localised with the centrosome and CD3ζ^+^ endosomes (Fig.1d, S2b). This result was confirmed in peripheral blood T cells purified from healthy donors and conjugated with Raji cells pulsed with a mix of SEA, SEB and SEE to cover a substantial proportion of the TCR Vβ repertoire and hence maximize the number of responding T cells (Fig.1e).

### BBS1 is required for centrosome translocation to the IS

Based on the homologies between the IS and the primary cilium [24–26], we hypothesized that BBS1, which is essential for ciliogenesis [41,42], could participate in IS assembly. To address this question we transduced Jurkat T cells with lentiviral particles containing an shRNA specific for BBS1 to generate a line stably knocked down for BBS1 expression (J KD; 90% depletion). A non-targeting shRNA was used to generate a control line (ctr) (Fig.S1b). The analysis was extended to primary T cells transiently knocked out for BBS1 expression (T KO; ~70% depletion) by CRISPR-Cas9 gene editing (Fig.S1b). BBS1 depletion did not affect either the levels of surface CD3 or expression of the other BBS core components (Fig.S1c,d).

The outcome of BBS1 depletion for IS assembly was assessed by confocal imaging using as readouts centrosome translocation to the IS and the synaptic accumulation of TCR:CD3 complexes in 15-min conjugates, a timepoint when IS maturation has fully occurred in ctr T cells. Additionally, signaling was assessed by quantifying tyrosine phosphoprotein accumulation at the IS in conjugates stained with anti-phosphotyrosine antibodies. BBS1 depletion in either Jurkat or primary T cells resulted in a defective accumulation of CD3ζ and tyrosine phosphoproteins at the IS (Fig.2a,b; Fig.S3a,b), concomitant with a failure to effectively polarize the centrosome (Fig.2a,b; Fig.S3a,b) and the endosomal TCR pool (Fig.2c) towards the APC.

**Fig. 2.**
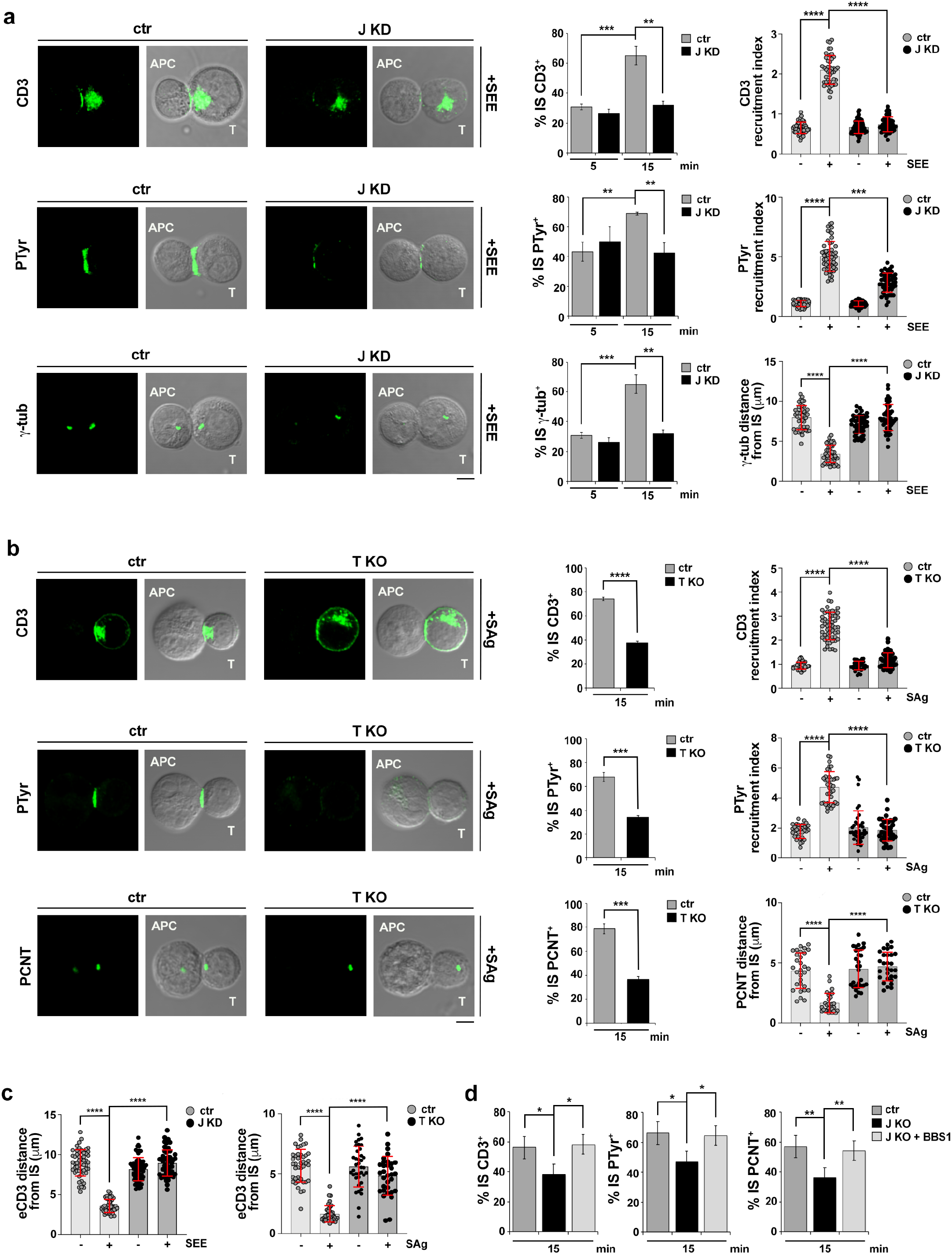
BBS1 is required for centrosome translocation to the IS. (**a**) *Left*, Immunofluorescence analysis of CD3ζ, tyrosine phosphoproteins (PTyr) and centrosome (γ-tubulin) in 15-min conjugates of control (ctr) and BBS1KD (J KD) Jurkat cells with SEE-loaded Raji B cells (APC). Representative images (medial optical sections) of the conjugates formed in the presence of SEE are shown (see Fig.S3a for representative images of conjugates formed in the absence of SEE). *Righ*t, Quantifications (%) of conjugates harboring CD3ζ, PTyr and γ-tubulin staining at the IS 5 min and 15 min after conjugate formation are shown on right of the images (≥ 25 cells/sample, n=3, unpaired two-tailed Student’s *t*-test). Top and middle histograms: Relative CD3ζ and PTyr fluorescence intensity at the IS in conjugates of control (ctr) and BBS1KD (J KD) Jurkat cells with Raji B cells in the presence or absence of SEE. The data are expressed as recruitment index, which is calculated as the *ratio* of CD3ζ and PTyr fluorescence intensity at the T cell:APC contact site to the total T cell area (≥ 10 cells/sample, n=3, Kruskal-Wallis test). *B*ottom histogram: Measurement of the distance (μm) of the centrosome (γ-tubulin) from the T cell:APC contact site in conjugates of control (ctr) and BBS1KD (J KD) Jurkat cells with Raji B cells in the presence or absence of SEE (≥ 10 cells/sample, n=3, Kruskal-Wallis test). (**b**) *Left*, Immunofluorescence analysis of CD3ζ, tyrosine phosphoproteins (PTyr) and centrosome (PCNT) in 15-min conjugates of control (ctr) and BBS1 KO (T KO) primary T cells with SAg-loaded (SEA+SEB+SEE) Raji B cells (APC). Representative images (medial optical sections) formed in the presence of SAg are shown (see Fig.S3b for representative images of conjugates formed in the absence of SAg). Quantifications (%) of conjugates harboring CD3ζ, PTyr and PCNT staining at the IS 15 min after conjugate formation are shown on right of the representative images (≥25 cells/sample, n=3, unpaired two-tailed Student’s *t*-test). *Right* (top and middle histograms), Relative CD3ζ and PTyr fluorescence intensity at the IS in 15-min conjugates of control (ctr) and BBS1KO (T KO) primary T cells with Raji B cells in the presence or absence of SAg. The data are expressed as recruitment index as in panel A (≥ 10 cells/sample, n=3, Kruskal-Wallis test). *Right* (bottom histogram), Measurement of the distance (μm) of the centrosome (PCNT) from the T cell:APC contact site in 15-min conjugates of control (ctr) and BBS1KO (T KO) primary T cells with Raji B cells in the presence or absence of SAg (≥ 10 cells/sample, n=3, Kruskal-Wallis test). (**c**) Measurement of the distance (μm) of the endosomal CD3ζ pool from the T cell:APC contact site in 15-min conjugates of control (ctr) and BBS1KD (J KD) Jurkat cells (*left*), or control (ctr) and BBS1 KO (T KO) primary T cells (*right*), with Raji B cells, in the presence or absence of SEE (*left*) or SAg (*right*) (≥ 10 cells/sample, n=3, Kruskal-Wallis test). (**d**) The histograms show the quantification (%) of conjugates with CD3ζ, PTyr or γ-tubulin staining at the IS 15 min after conjugate formation of control (ctr) or BBS1 KO (J KO) Jurkat cells, the latter transfected with either empty vector (ctr, J KO) or the same vector encoding wild-type BBS1 (J KO+BBS1-GFP), with SEE-loaded Raji B cells (≥ 25 cells/sample, n=3, unpaired two-tailed Student’s *t*-test). Size bar, 5 μm. The data are expressed as mean±SD. ****P≤0.0001; ***P≤0.001; **P≤0.01; *P≤0.05.

To understand whether the IS formation defects in BBS1 KD cells were causally linked to BBS1 deficiency we generated a BBS1 knockout Jurkat line (J KO) by CRISPR-Cas9 gene editing (Fig.S1b,c). These cells recapitulated the IS defects observed in BBS1 KD cells (Fig.2d). Restoring BBS1 expression in BBS1 KO cells by transfection with a BBS1-GFP encoding construct (Fig.S1e,f) rescued the ability of these cells to promote centrosome polarization and synaptic CD3ζ and tyrosine phosphoprotein accumulation (Fig.2d). Hence BBS1 is required the the assembly of functional T cell synapses.

Centrosome translocation is triggered in response to TCR recognition of cognate pMHC. This allows for the polarized transport to the IS not only of endosomal TCRs, but also of key signaling mediators associated with recycling endosomes, including the initiating kinase Lck [2], the transmembrane adaptor LAT [4] and the small GTPase Rac1 [19], all of which contribute to sustain signaling during T cell activation. To understand whether the defect in CD3 accumulation and phosphotyrosine signaling at the IS formed by BBS1-deficient T cells in 15-min conjugates occurs early during IS assembly or downstream of centrosome polarization we measured these events at an earlier time point (5 min). Normal CD3 and tyrosine phosphoprotein enrichment at the T cell interface with the APC could be observed in 5-min antigen-specific conjugates formed by BBS1 KD Jurkat cells (Fig.2a), suggesting that the signaling events triggered by the membrane-associated TCR:CD3 pool that eventually lead to centrosome translocation and polarized TCR recycling are not affected by BBS1 deficiency. These results were confirmed by flow cytometric analysis of protein tyrosine phosphorylation in T cells activated at different time points with SEE-pulsed APCs (Fig.S3c). Hence BBS1 participates in IS assembly by promoting the polarization of the centrosome to the IS, which in turn allows for the local recruitment of TCRs associated with recycling endosomes.

### BBS1 is required for centrosome untethering from the nucleus during IS formation

In quiescent cells the centrosome is tethered to the nucleus by a cytoskeletal network [43]. The LInkers of the Nucleoskeleton and Cytoskeleton (LINC) protein complexes are essential to keep the centrosome attached to the nucleus by bridging centrosome-nucleated microtubules and actin filaments (F-actin) to the nuclear envelope [43,44]. In resting B cells, the LINC complex component nesprin-2 and F-actin tether the centrosome to the nucleus [37]. F-actin clearance at the centrosome due to a reduction in F-actin nucleation allows for centrosome detachment and its translocation to the B cell IS [37].

We hypothesized that the inability of the centrosome to translocate to the IS in BBS1-deficient T cells may reflect a role for BBS1 in the regulation of centrosome tethering to the nucleus. To test this possibility we first measured the distance of the centrosome from the nuclear membrane in SEE-specific conjugates of control and BBS1 KD Jurkat cells co-stained for the centrosomal markers pericentrin (PCNT) or PCM-1 and the LINC complex component nesprin-2. Conjugates formed in the absence of SEE were used as controls. The distance between the centrosome and the nuclear membrane increased in antigen-specific conjugates of ctr cells, concomitant with centrosome translocation to the IS (Fig.3a; Fig.S5a-c). In contrast, no difference in the distance of the centrosome from the nuclear membrane could be observed in conjugates formed by BBS1 KD Jurkat cells in the absence or presence of SEE (Fig.3a; Fig.S5c). These observations were validated in primary BBS1 KO T cells (Fig.3b). Hence BBS1 is required for centrosome untethering from the nucleus, accounting for the defect in centrosome translocation to the IS observed in BBS1-deficient T cells.

**Fig. 3.**
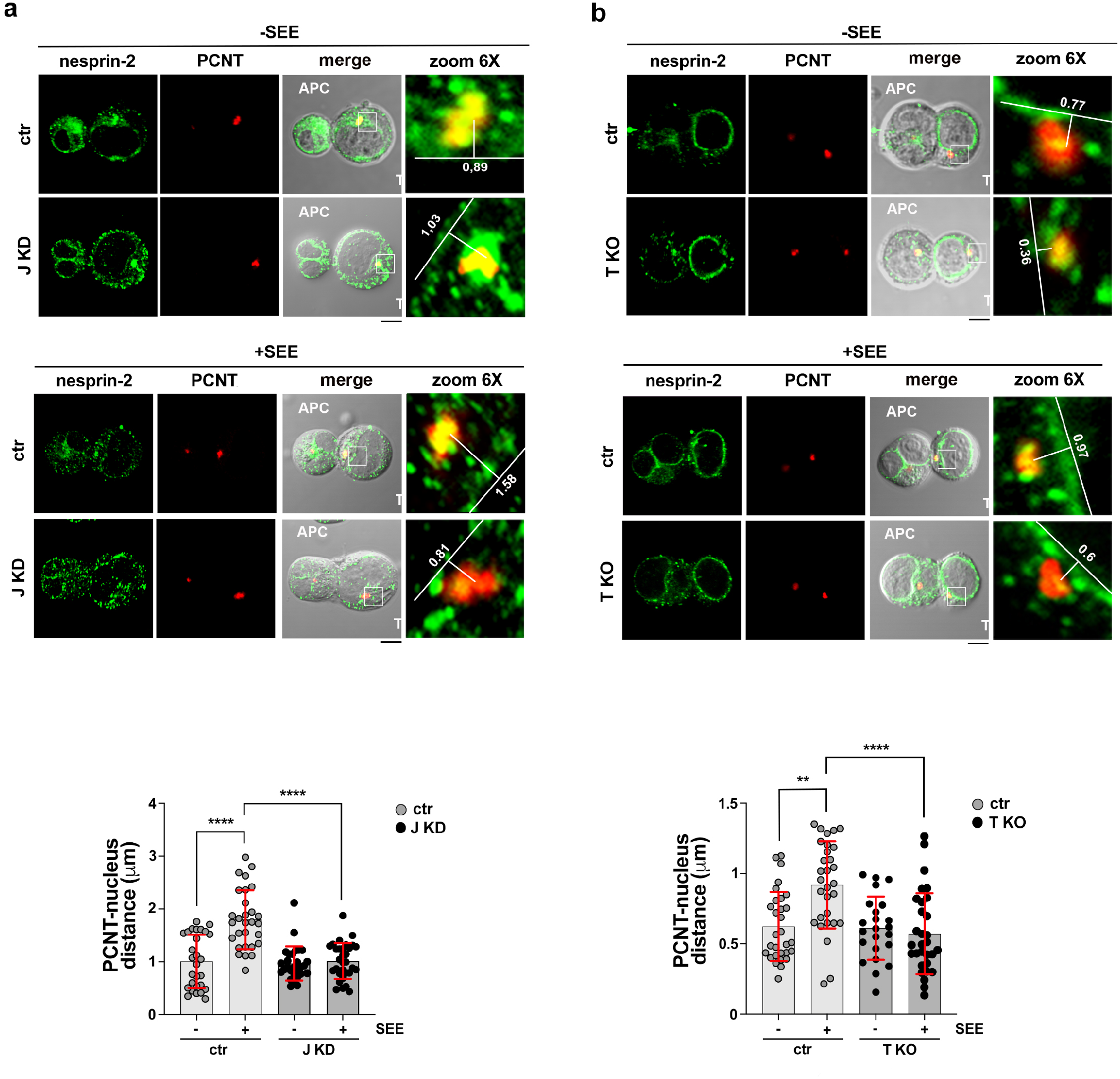
BBS1 is required for centrosome untethering from the nucleus during IS formation. (**a,b**) *Top*, Immunofluorescence analysis of the distance between the centrosome (PCNT) and the nuclear membrane (nesprin-2) in 15-min conjugates of control (ctr) and BBS1KD (J KD) Jurkat cells (**a**), or control (ctr) and BBS1 KO (T KO) primary T cells (**b**) and Raji B cells (APC) formed in the presence or absence of SEE (Jurkat cells) or SAg (T cells). The centrosomal area was magnified to highlight the parameters used for quantification (detailed in Fig.S4a). *Bottom*, Quantifications of the centrosome-nucleus distance (μm) in conjugates of which representative images (medial optical sections) are shown in (**a,b**) (≥ 10 cells/sample, n=3, Kruskal-Wallis test). Size bar, 5 μm. The data are expressed as mean±SD. ****P≤0.0001; **P≤0.01.

### BBS1 is required for centrosomal F-actin clearance during IS formation

Centrosomal F-actin tethers the centrosome to the nucleus by interacting with nesprins, which cross the outer nuclear membrane and interact with the SUN proteins resident in the nuclear envelope lumen [45]. Similar to B cells [37], F-actin clearance from the centrosome has been recently shown to occur during IS formation in T cells, allowing for centrosome detachment and polarization [46]. Conjugate staining with fluorochrome-labelled phalloidin and anti-γ-tubulin antibodies revealed the presence of a centrosomal F-actin pool surrounded by a radial pattern of F-actin^+^ dots (Fig.4a).

**Fig. 4.**
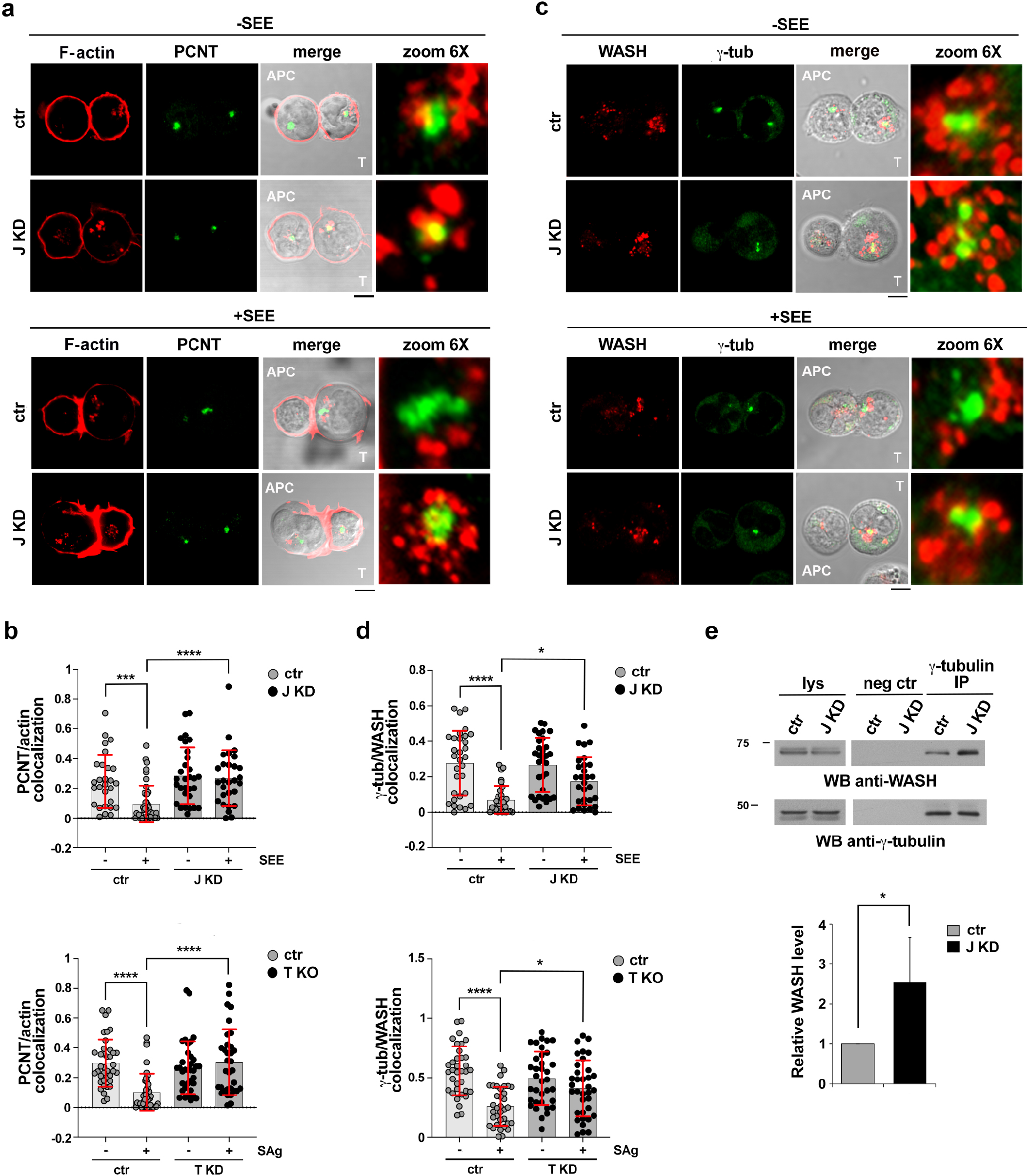
BBS1 is required for centrosomal F-actin clearance during IS formation. (**a**) Immunofluorescence analysis of centrosomal F-actin (phalloidin) in 15-min conjugates of control (ctr) and BBS1KD (J KD) Jurkat cells with Raji B cells (APC) in the absence or presence of SEE, co-stained for PCNT. (**b**) Quantification using Mander’s coefficient of the weighted colocalization of PCNT with centrosomal F-actin. The parameters used for quantification as detailed in Fig.S4b. The histograms show the quantification in 15-min conjugates of control (ctr) and BBS1KD (J KD) Jurkat cells with Raji B cells (APC) in the absence or presence of SEE (*top*), or conjugates of control (ctr) and BBS1 KO (T KO) primary T cells in the absence or presence of SAg (*bottom*) (≥ 10 cells/sample, n=3, Kruskal-Wallis test). (**c**) Immunofluorescence analysis of centrosomal WASH in 15-min conjugates of control (ctr) and BBS1KD (J KD) Jurkat cells with Raji B cells (APC) in the absence or presence of SEE, co-stained for γ-tubulin. (**d**) Quantification using Mander’s coefficient of the weighted colocalization of γ-tubulin with centrosomal WASH. The parameters used for quantification as detailed in Fig.S4b. The histograms show the quantification in 15-min conjugates of control (ctr) and BBS1KD (J KD) Jurkat cells with Raji B cells (APC) in the absence or presence of SEE (*top*) or conjugates of control (ctr) and BBS1 KO (TKO) primary T cells in the absence or presence of SAg (*bottom*) (≥ 10 cells/sample, n=3, Kruskal-Wallis test). Representative images (medial optical sections) are shown. Size bar, 5 μm. (**e**) Immunoblot analysis of γ-tubulin-specific immunoprecipitates from lysates of control (ctr) and BBS1KD (J KD) Jurkat cells using anti-WASH antibodies. A preclearing control (proteins that bound to Protein-A-Sepharose before the addition of primary antibody) is shown (neg ctr). Total cell lysates (lys) were included in each gel to identify the migration of the tested proteins. The migration of molecular mass markers is shown for each filter. The histogram shows the relative intensities of the immunoreactive bands corresponding to dynein (J KD vs ctr) normalized to the immunoprecipitated γ-tubulin (mean±SD, n = 3, ctr value = 1, one-sample *t*-test). ****P≤0.0001; ***P≤0.001.

A colocalization analysis showed that these corresponded to early endosomes, where actin is known to polymerize locally to promote recycling [47], and to a lesser extent, to recycling endosomes (Fig.S6a,b). Measuring the co-localization of F-actin with PCNT showed a decrease in centrosomal F-actin in SEE-specific conjugates of control cells compared to conjugates formed in the absence of SEE. Conversely, the co-localization of F-actin with the centrosome did not change in antigen-specific conjugates formed by BBS1 KD Jurkat cells (Fig.4a; Fig.S5d). Similar results were obtained using primary control and BBS1 KO T cells (Fig.4b).

The actin regulator WASH has been shown to promote F-actin accumulation at the centrosome by recruiting the branched F-actin nucleator Arp2/3 [48,49,37]. Co-localization analyses in control conjugates stained with anti-WASH and anti-γ-tubulin antibodies revealed a centrosomal WASH pool that decreased in antigen-specific conjugates formed by control T cells, but not by BBS1-deficient T cells (Fig.4c,d). Interestingly, more WASH co-immunoprecipitated with γ-tubulin in BBS1 KD cells compared to controls (Fig.4e), suggesting that WASH association with the centrosome is negatively regulated by BBS1. Collectively, these results indicate that BBS1 deficiency results in the failure of T cells to clear centrosomal F-actin, at least in part due to a higher stability of the centrosomal WASH pool, which likely accounts for the inability of the centrosome to disengage from the nucleus and translocate to the IS.

### Centrosome translocation to the IS is regulated by the proteasome

The composition and function of the centrosome is dynamically regulated by a local proteasome pool that fine-tunes the concentration of cell cycle regulators, cell fate determinants and other mediators of key cellular processes in a variety of cell types, including immune cells [50]. Interestingly, proteasome activity has been recently shown to be essential for F-actin remodeling at the centrosome to allow for its disengagement from the nucleus and translocation to the IS in B cells [51].

To assess the role of the proteasome in F-actin clearance from the centrosome and its translocation towards the T cell IS we investigated the impact of proteasome inhibitors on the ability of T cells to form functional immune synapses. Jurkat or primary T cells were treated for 2 h with two different proteasome inhibitors, MG132 or epoxomicin, prior to conjugate formation. These pre-treatments resulted in the expected accumulation of ubiquitylated proteins, as assessed by immunoblot analysis of cell lysates (Fig.S7a), without any significant loss in cell viability (Fig.S7b). Remarkably, proteasome inhibitor pre-treatment phenocopied the effects of BBS1 deficiency on IS assembly, with a defect in centrosome polarization associated with its inability to untether from the nuclear envelope, as assessed by conjugate co-staining with anti-PCNT and anti-nesprin-2 antibodies (Fig.5a,b,c,d). Additionally, the endosomal TCR pool failed to translocate to the IS, concomitant with an impaired synaptic accumulation of TCR and tyrosine phosphoproteins (Fig.5e,f,g,h). Similar to BBS1-deficient T cells, the centrosomal F-actin and WASH pools did not become depleted during IS formation in Jurkat (Fig.5i) or primary (Fig.5j) T cells treated with proteasome inhibitors (Fig.5i-j). Although proteasome inhibition may result in alterations in other ubiquitin-regulated processes, these results suggest a functional link between the ubiquitin-proteasome system and the F-actin clearance-related centrosome detachment from the nucleus, on which its translocation towards the T cell:APC interface and the assembly of a functional IS crucially depend.

**Fig. 5.**
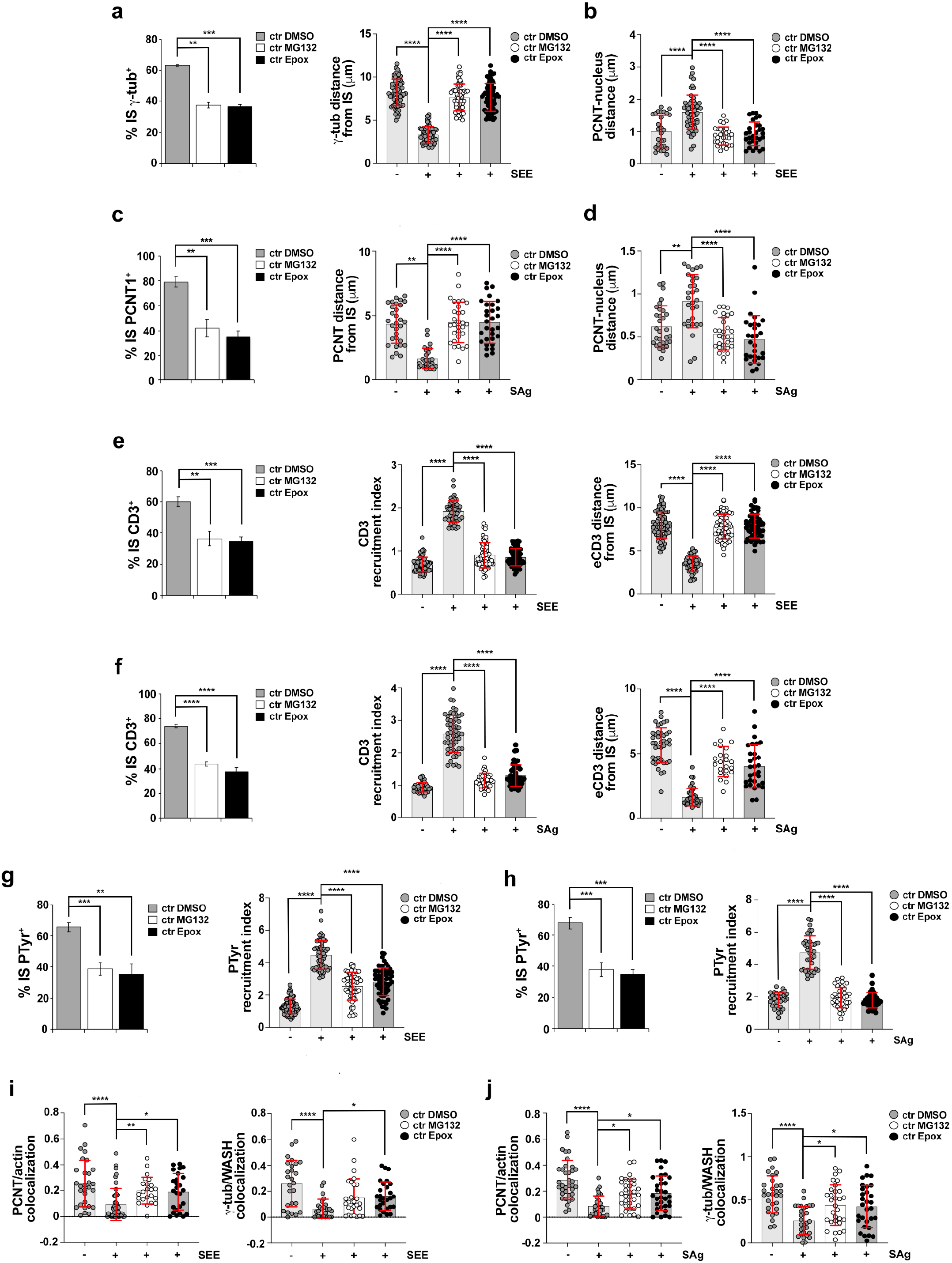
Centrosome translocation to the IS is regulated by the proteasome. (**a,c**) *Left*, Quantifications (%) of 15-min conjugates harboring centrosome (γ-tubulin in Jurkat cells, PCNT in primary T cells) staining in conjugates of control (ctr) Jurkat (**a**) or primary (**c**) T cells, either untreated (DMSO) or pre-treated with the proteasome inhibitors MG132 or epoxomicin (Epox) (≥25 cells/sample, n=3, unpaired two-tailed Student’s *t*-test). *Right*, Measurement of the distance (μm) of the centrosome (γ-tubulin in Jurkat cells, PCNT in primary T cells) from the T cell:APC contact site in 15-min conjugates of control (ctr) Jurkat (**a**) or primary (**c**) T cells. Conjugates formed in the absence of SEE (**a**) or SAg (**c**) were used as negative controls (≥ 10 cells/sample, n=3, Kruskal-Wallis test). (**b,d**) Measurement of the centrosome-nucleus distance (μm) in 15-min SEE/SAg-specific conjugates of control (ctr) Jurkat (**b**) or primary (**d**) T cells, either untreated (DMSO) or pre-treated with the proteasome inhibitors MG132 or epoxomicin. Conjugates formed in the absence of SEE (**b**) or Sag (**d**) were used as negative controls (≥ 10 cells/sample, n=3, Kruskal-Wallis test). The parameters used for quantification are detailed in Fig.S4a. (**e,f**) *Left*, Quantifications (%) of 15-min conjugates harboring CD3ζ staining in SEE/SAg-specific conjugates of control (ctr) Jurkat (**e**) or primary (**f**) T cells, either untreated (DMSO) or pre-treated with the proteasome inhibitors MG132 or epoxomicin (≥25 cells/sample, n=3, unpaired two-tailed Student’s *t*-test). *Middle*, Relative CD3ζ fluorescence intensity at the IS in 15-min SEE/SAg-specific conjugates of control (ctr) Jurkat (**e**) or primary (**f**) T cells. Conjugates formed in the absence of SEE (**e**) or SAg (**f**) were used as negative controls (≥ 10 cells/sample, n=3, Kruskal-Wallis test). The data are expressed as recruitment index, which is calculated as the *ratio* of CD3ζ fluorescence intensity at the T cell:APC contact site to the total T cell area. *Right,* Measurement of the distance (μm) of the endosomal TCR/CD3 pool from the T cell:APC contact site in 15-min SEE/SAg-specific conjugates of control (ctr) Jurkat (**e**) or primary (**f**) T cells. Conjugates formed in the absence of SEE/SAg were used as negative controls (≥ 10 cells/sample, n=3, Kruskal-Wallis test). (**g,h**) *Top*, Quantifications (%) of conjugates harboring PTyr staining at the IS in 15-min SEE/SAg-specific conjugates of control (ctr) Jurkat (**g**) or primary (**h**) T cells, either untreated (DMSO) or pre-treated with the proteasome inhibitors MG132 or epoxomicin (≥ 25 cells/sample, n=3, unpaired two-tailed Student’s *t*-test). *Bottom*, Relative PTyr fluorescence intensity at the IS in SEE/SAg-specific conjugates of control (ctr) Jurkat (**g**) or primary (**h**) T cells. Conjugates formed in the absence of SEE (**g**) or SAg (**h**) were used as negative controls (≥ 10 cells/sample, n=3, Kruskal-Wallis test). The data are expressed as recruitment index, which is calculated as the *ratio* of PTyr fluorescence intensity at the T cell:APC contact site to the total T cell area. (**i,j**) Quantification using Mander’s coefficient of the weighted colocalization of PCNT with centrosomal F-actin (*left*) or γ-tubulin with centrosomal WASH (*right*) is shown. The parameters used for quantification are detailed in Fig.S4B. The histograms show the quantification in SEE/SAg-specific conjugates of control (ctr) Jurkat (**i**) or primary (**j**) T cells, either untreated (DMSO) or pre-treated with the proteasome inhibitors MG132 or epoxomicin. Conjugates formed in the absence of SEE (**i**) or Sag (**j**) were used as negative controls (≥ 10 cells/sample, n=3, Kruskal-Wallis test). The data are expressed as mean±SD. ****P≤0.0001; ***P≤0.001; **P≤0.01; *P≤0.05.

### The 19S regulatory subunit of the proteasome is recruited to the centrosome during IS assembly through BBS1-mediated coupling to dynein

The 26S proteasome consists of a 20S catalytic subunit and a 19S regulatory subunit (19S RP) that is essential not only for its proteolytic activity but also for substrate recognition, unfolding and translocation into the 20S internal core [52]. We hypothesized that the activity of the centrosomal proteasome could be regulated by the dynamic recruitment of its regulatory subunit during IS assembly to promote local F-actin clearance. To address this question, we imaged the 19S RP pool associated with the centrosome at different time points after conjugate formation. A centrosomal accumulation of 19S RP was detected in control T cells under basal conditions, as assessed in conjugates formed in the absence of antigen and co-stained for γ-tubulin (Fig. 6a). This pool increased early during IS formation (1 min), prior to the stabilization of the IS to its mature architecture featuring the centrosome polarized towards the APC (15 min) (Fig.6a). Interestingly, a reduction in centrosomal 19S RP was observed under the same conditions in BBS1-deficient T cells compared to control cells (Fig.6a; Fig.S5e). These results suggest that the activity of the centrosome-associated proteasome is upregulated during IS formation through 19S RP recruitment to allow for local F-actin clearance and centrosome detachment from the nucleus. In agreement with this hypothesis, co-staining antigen-specific conjugates with anti-ubiquitin and anti-PCNT antibodies revealed a centrosomal accumulation of ubiquitylated proteins in BBS1-deficient T cells (Fig.6b; Fig.S5f) in the absence of a generalized defect in proteasome activity, as assessed by probing post-nuclear supernatants with anti-ubiquitin antibodies (Fig.S7c).

**Fig. 6.**
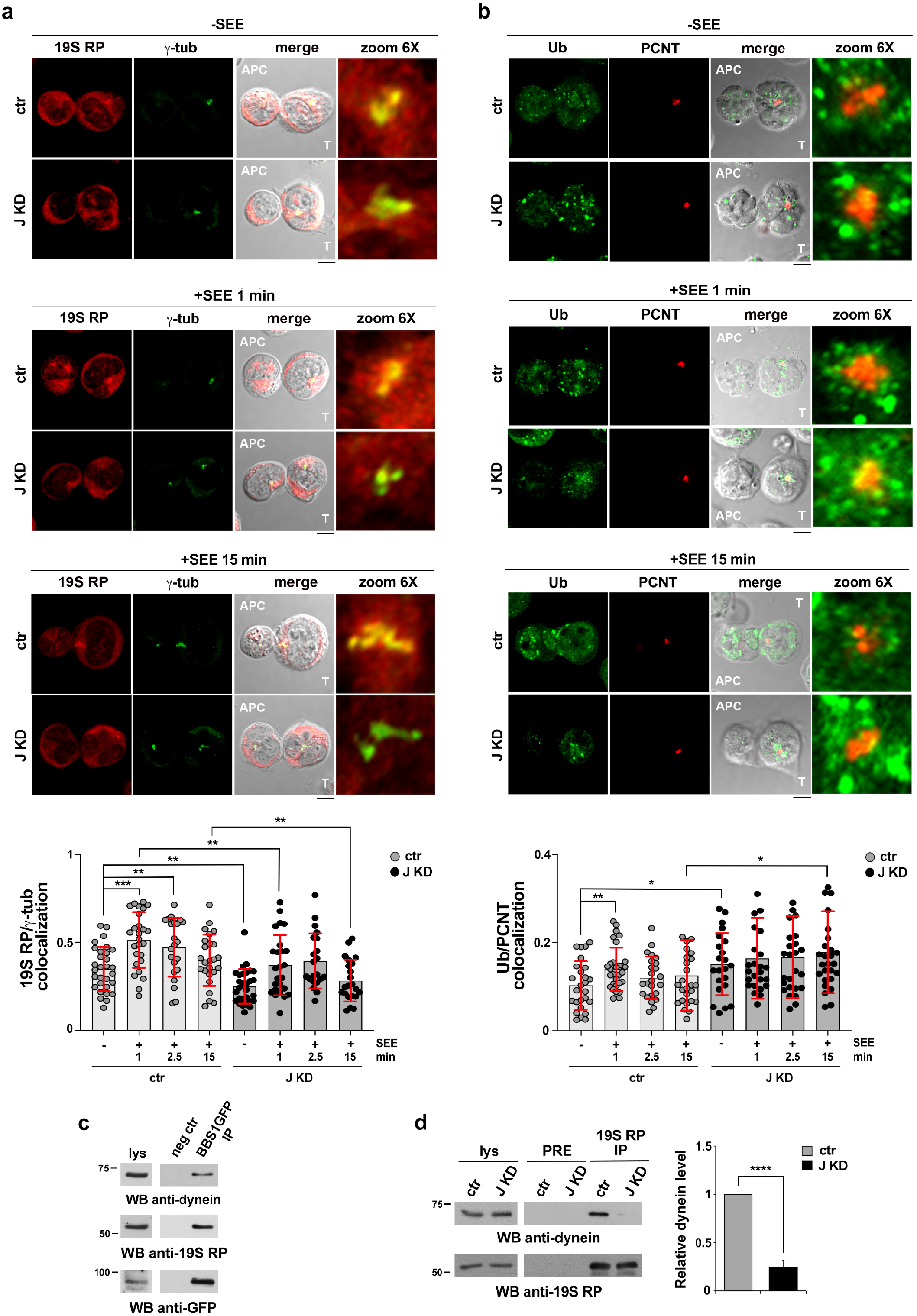
The 19S regulatory subunit of the proteasome is recruited to the centrosome during IS assembly through BBS1-mediated coupling to dynein. (**a**) Time-course immunofluorescence analysis of 19S RP recruitment to the centrosome (γ-tubulin) in conjugates of control (ctr) and BBS1KD (J KD) Jurkat cells with SEE-loaded Raji B cells (1 min and 15 min) or with Raji B cells in the absence of SEE (1 min). Quantification using Mander’s coefficient of the weighted colocalization of 19S RP with the centrosome (γ-tubulin) is shown in the bottom panel (≥ 10 cells/sample, n=3, Mann-Whitney test). The parameters used for quantification are detailed in Fig.S4b. (**b**) Time-course immunofluorescence analysis of ubiquitin (Ub) recruitment to the centrosome (PCNT) in conjugates of control (ctr) and BBS1KD (J KD) Jurkat cells with SEE-loaded Raji B cells (1 min and 15 min) or with Raji B cells in the absence of SEE (1 min). Quantification using Mander’s coefficient of the weighted colocalization of ubiquitin with the centrosome (PCNT) is shown in the bottom panel (≥ 10 cells/sample, n=3, Mann-Whitney test). The parameters used for quantification are detailed in Fig.S4b. Representative images (medial optical sections) are shown. Size bar, 5 μm. (**c**) Immunoblot analysis of GFP-specific immunoprecipitates (BBS1-GFP IP) from lysates of BBS1-GFP-expressing cells (mean±SD, n=3). A preclearing control (proteins that bound to Protein-A-Sepharose before the addition of primary antibody) is shown (neg ctr). A total cell lysate (lys) was included in each gel to identify the migration of the tested proteins. The migration of molecular mass markers is shown for each filter. (**d**) Immunoblot analysis of 19S RP-specific immunoprecipitates from lysates of control (ctr) and BBS1KD (J KD) Jurkat cells. A preclearing control (proteins that bound to Protein-A-Sepharose before the addition of primary antibody) is shown (neg ctr). Total cell lysates (lys) were included in each gel to identify the migration of the tested proteins. The migration of molecular mass markers is shown for each filter. The histogram shows the relative intensities of the immunoreactive bands corresponding to dynein (intermediate chain) (J KD vs ctr) normalized to the immunoprecipitated 19S RP (mean±SD, n≥3, ctr value = 1, one-sample *t*-test). ****P≤0.0001; ***P≤0.001; **P≤0.01; *P≤0.05.

Microtubule-dependent retrograde transport, involving coupling to the minus-end microtubule motor dynein, has been implicated in the regulation of local proteasome pools during axon development [53]. A similar process may account for the centrosomal accumulation of 19S RP. To address this issue we assessed the ability of BBS1 to interact with 19S RP in co-immunoprecipitation assays on a Jurkat transfectant generated using GFP-tagged BBS1. BBS1 was found to constitutively associate with 19S RP (Fig.6c). Consistent with the role of the BBSome in mediating cargo-dynein interactions during retrograde transport [28], dynein (intermediate chain) was also found to co-immunoprecipitate with BBS1-GFP (Fig.6c). BBS1 deficiency led to a disruption of the interaction between 19S RP and dynein (Fig.6d). Together, these results suggest that BBS1 promotes centrosome translocation during IS assembly by assisting the dynein-dependent recruitment of 19S RP to the centrosome to allow for the proteasome-regulated clearance of centrosomal F-actin, leading to centrosome detachment from the nucleus.

## DISCUSSION

Based on rapidly accumulating evidence that proteins implicated in the growth and function of the primary cilium are co-opted by non-ciliated cells to regulate basic processes, vesicular trafficking being a prominent example thereof [7], here we have explored the role of the BBSome component BBS1 in the formation of the T cell IS. We show that BBS1 participates in IS assembly by promoting centrosome polarization towards the APC and that defective centrosome translocation in BBS1-deficient T cells results from its failure to efficiently detach from the nuclear envelope, concomitant with defective centrosomal clearance of F-actin and its regulator WASH. The observation that proteasome inhibitors phenocopy the defects caused by BBS1 deficiency suggested a link between centrosome untethering form the nucleus and activity of the centrosome-associated proteasome, involving BBS1. Consistent with this notion, we found that BBS1 couples the 19S regulatory subunit of the proteasome to dynein for its transport to the centrosome in the early stages of IS formation (Fig.7). These results identify BBS1 as a key player in IS assembly.

BBS1 is responsible for binding BBS3/Arl6 to promote BBSome entry into cilia [29]. Additionally, assembly of a pre-BBSome has been recently reported in a preliminary form to be primed by BBS4 at pericentriolar satellites, with basal body-resident BBS1 targeting the pre-BBSome to the base of the cilium to locally complete BBSome maturation [38]. Given this pivotal role of BBS1 in BBSome assembly, we believe that our results obtained on BBS1 reflect the function of the entire BBS complex, as shown in ciliated cells [54–56]. We cannot however rule out a BBSome-independent function of BBS1 in centrosome untethering from the nucleus for IS polarization.

**Fig. 7.**
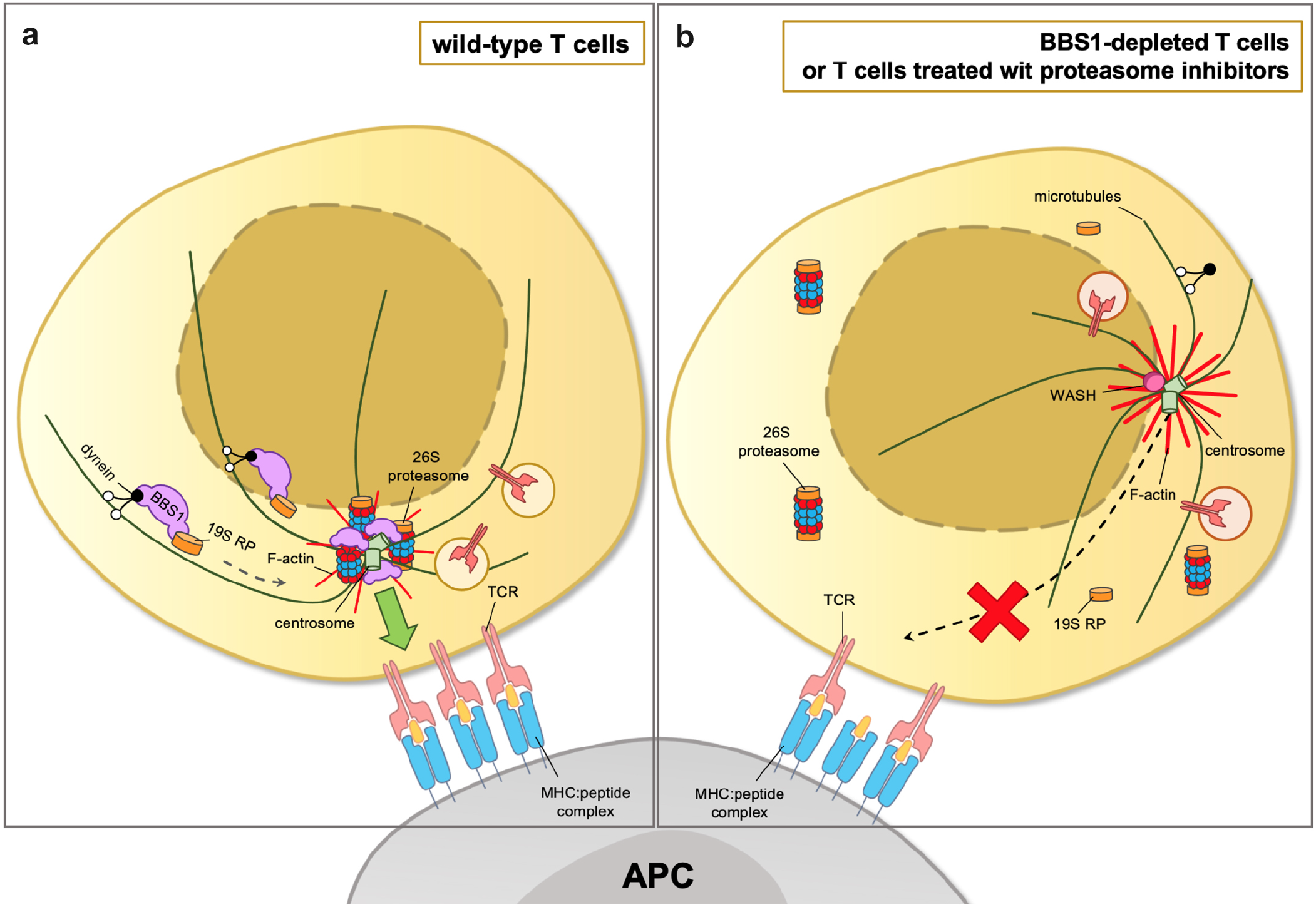
Proposed model of regulation of centrosome polarization by BBS1 during IS assembly in T cells. (**a**) Centrosome polarization during IS formation is paralleled by a rapid change in F-actin dynamics at the centrosomal area towards decreased actin nucleation that favors centrosome detachment from the nucleus. A centrosome-associated proteasome pool promotes clearance of centrosomal F-actin and the actin nucleator factor WASH, allowing for centrosome separation from the nucleus. This process is enabled by BBS1, which promotes the dynein-dependent recruitment of the 19S regulatory subunit (19S RP) to the centrosome. (**b**) When T cells are depleted of BBS1 or treated with proteasome inhibitors, they fail to promote the proteasome-dependent clearance of centrosomal F-actin. This results in the inability of the centrosome to untether from the nucleus and translocate to the IS, leading to impaired cell polarity and a defective formation of functional immune synapses.

The role of the BBSome in ciliary protein trafficking has been extensively documented. While its implication in protein entry into the cilium is as yet debated, the main function of the BBSome is to cooperate with the IFT-B complex in retrograde trafficking of ciliary receptors to the base of the cilium and their crossing of the transition zone before returning to the cell body [27,28]. This trafficking-related function of the BBSome, which may result from its ability to polymerize a flat coat on synthetic membranes [57], has been reported for a number of receptors, GPCRs being the main group [58,59]. Our results highlight an apparent divergence in the function of the BBSome in ciliated cells and in the non-ciliated T cells, since in the latter BBS1 is required for the transport of a cytosolic protein complex, the 19S proteasome subunit. However, since centrosome translocation is a prerequisite for reorientation of the secretory apparatus towards the IS and focalized exocytosis [60,61], we cannot exclude that the BBSome is also implicated in trafficking events downstream of centrosome translocation during IS formation, in particular TCR recycling, in which IFT20 and other IFT-B complex components participate [13,14].

The centrosome has been shown to promote the local polymerization of actin filaments through the local recruitment of Arp2/3 and one of its activators, WASH [49]. A central function of this F-actin is to anchor the centrosome to the nucleus via their interaction with the LINC complex [43]. These interactions must be disrupted when the centrosome has to move to a different subcellular localization, such as during IS formation. During IS assembly the centrosomal F-actin pool undergoes depletion [37,46]. In B cells this has been demonstrated to be due to a relocalization of Arp2/3 from the centrosome to the IS, which leads to centrosome detachment from the nucleus and allows for its polarization [37]. Here we show that centrosome F-actin clearance is impaired in BBS1-deficient T cells undergoing IS assembly concomitant with the persistance of a centrosomal pool of WASH, which results in the failure of the centrosome to untether from the nuclear membrane to translocate towards the T cell:APC contact. Interestingly, similar to B cells [51], we found that centrosomal F-actin clearance is controlled by proteasome activity in T cells. The impairment in 19S RP recruitment to the centrosome of BBS1-deficient T cells at early stages of IS assembly, when the centrosome undergoes polarization to the nascent IS, paralleled by a centrosomal accumulation of ubiquitylated proteins, supports the notion that BBS1 modulates the centrosome proteasome. To carry out this function it exploits its documented cilia-related retrograde trafficking function to couple the 19S regulatory subunit of the proteasome to dynein. It is noteworthy that BBS proteins have been shown to interact with proteasome subunits in ciliated cells, and that their loss led to proteasome depletion from the centrosome [54]. Although the underlying mechanism has not been investigated, these data suggest a conserved, cilia-independent function of BBS proteins in centrosome proteostasis in ciliated and non-ciliated cells. Interestingly, the BBSome also interacts with ubiquitin to promote the removal of Smoothened and other GPCRs from the ciliary membrane [62–64]. This function might be relevant for the fate of post-endocytic synaptic TCRs.

The molecular targets of the centrosomal proteasome that undergo degradation to allow for local F-actin clearance remain to be elucidated. Our finding that the centrosomal WASH pool becomes depleted during IS formation highlights WASH as a possible candidate. Although to date there is no published evidence of a direct proteasomal regulation of WASH itself, the activity of WASH has been reported to be indirectly regulated by the proteasome, which degrades the E3 ubiquitin ligase TRIM27 that activates WASH by non-degradative K63-linked ubiquitylation [65]. Additionally, several proteins that promote F-actin nucleation by binding the Arp2/3 complex are known to be regulated by proteasome-mediated degradation, including WASp [66] and WAVE2 [67]. Another potential proteasome-regulated target is the centriolar satellite protein PCM-1 [68], which contributes to the actin-nucleating function of the centrosome by recruiting WASH and Arp2/3 [49]. Degradation of any of these proteins by the centrosome-associated proteasome, the activity of which is indirectly regulated by BBS1 via dynein-dependent transport of 19S RP, is expected to slow down the rate of F-actin polymerization, leading to a shift of actin dynamics towards turnover.

BBS is a pleiotropic genetic disorder characterized by a number of abnormalities that include polydactyly, kidney dysfunction, obesity, hypogonadism and anosmia [69,70]. These defects are caused by dysfunction of primary cilia which in general grow normally but are unable to signal due to impaired BBSome-mediated ciliary trafficking of components of key signaling pathways, such as the Shh pathway [71]. The potential impact of BBS-related mutations on the immune system had not been addressed until recently. Tsyklauri and colleagues have now reported immune abnormalities in BBS patients, which are associated with a higher incidence of autoimmune diseases [72]. Using the *bbs4*^−/−^ mouse model of BBS, they showed that BBS4 deficiency affects mainly the B-cell compartment through a non-intrinsic mechanism that remains to be defined, with only minor effects on the T-cell compartment. However, they did not observe any significant effect of BBS4 deficiency on mouse B-cell and T-cell responses [72]. Our results, implicating BBS1 in IS assembly, are in apparent contrast with these findings. We can propose several factors that could underlie these differences. First, we have used human T cells, and emerging evidence has highlighted features of the human immune response that are not recapitulated in the mouse system [73,74]. Second, we have genetically inactivated *bbs1*, while Tsyklauri et al have targeted *bbs4*. The proteins encoded by these genes carry out complementary yet different functions in BBSome assembly [38]. Additionally, we cannot rule out at this stage BBSome-independent functions of BBS1 and BBS4 in T cells or other non-ciliated cells. Third, the fact that no defects in conjugate formation have been observed in mouse *bbs4*^−/−^ T cells [72] suggests that T cell adhesion to APCs is normal but does not exclude defects in IS assembly. Finally, compensatory mechanisms may account for the normal in vivo *bbs4^−/−^* T-cell response in the diabetes mouse model used by the authors [72].

The study of T cells from BBS patients will shed light on the outcome of BBSome dysfunction on T-cell IS formation and on the downstream events. Nonetheless, the fact that all BBSome components are expressed in T cells, the preliminary evidence provided in the report by Tsyklauri et al [72] and the data presented in this report, open a new perspective in the search for gene defects in primary immunodeficiencies of unknown aetiology.

## Acknowledgments

We wish to thank Max Nachury, Valentina Cianfanelli and Federico Galvagni for reagents and helpful suggestions, Claire Hivroz for critical reading of the manuscript and Maria Isabel Yuseff for useful advice.

## Abbreviations

APC: Antigen Presenting Cell
Arl: ADP Ribosylation Factor Like
Arp2/3: Actin-Related Protein 2/3 Complex
BBS: Bardet-Biedl syndrome
CRISPR: Clustered Regularly Interspaced Short Palindromic Repeats
EB-1: End-Binding 1
F-actin: Filamentous actin
FIP3: Rab11 Family-Interacting Protein 3
GFP: Green Fluorescent Protein
GMAP210: Golgi Microtubule-Associated Protein 210
GPCR: G Protein-coupled Receptor
IFT: IntraFlagellar Transport
IS: Immune Synapse
KD: Knock Down
KO: Knock Out
LAT: Linker for Activation of T cells
LINC: LInkers of the Nucleoskeleton and Cytoskeleton
PAS: Protein A Sepharose
PCM-1: PeriCentriolar Material-1
PCNT: Pericentrin
RT-PCR: Reverse Transcription Polymerase Chain Reaction
sgRNAs: small guide RNAs
SAg: SuperAntigen
SEA: Staphylococcal Enterotoxin A
SEB: Staphylococcal Enterotoxin B
SEE: Staphylococcal Enterotoxin E
SNARE: SNAP Receptor
Shh: Sonic hedgehog
TCR: T Cell Receptor
Unc-119: Uncoordinated-119
VAMP: Vesicle Associated Membrane Protein
WASH: WASP & Scar Homologue
WASp: Wiskott–Aldrich Syndrome protein
WAVE: WASP family Verprolin-homologous protein
19S RP: 19S proteasome Regulatory Subunit

## Declarations

### Funding

This work was carried out with the support of Fondazione Telethon, Italy (Grant GGP16003) to CTB. The support of AIRC (IG-2017 - ID 20148) to CTB is also gratefully acknowledged.

### Competing interests

The authors declare no financial or competing interest.

### Ethics approval

Buffy coats from anonymous healthy donors obtained from the Siena University Hospital blood bank were used for this study. The study was approved by the local ethics committee (Siena University Hospital)

### Consent to participate

Informed consent was obtained from blood donors by the physician in charge of the Siena University Hospital blood bank. Buffy coats were anonymised prior to distribution.

### Consent for publication

N/A

### Availability of data and material

All unique/stable reagents generated in this study are available from the Lead Contact without restriction. The datasets supporting the current study have not been deposited in a public repository but are available from the corresponding author on reasonable request.

### Code availability

N/A

### Author contributions

All authors contributed to the study conception and design. Material preparation, data collection and analysis were performed by Chiara Cassioli, Anna Onnis, Francesca Finetti, Nagaja Capitani and Ewood B Compeer. The first draft of the manuscript was written by Chiara Cassioli, Anna Onnis and Cosima T Baldari. All authors commented on previous versions of the manuscript. All authors read and approved the final manuscript.

## Notes

### Competing Interest Statement

The authors have declared no competing interest.

## REFERENCES

1. Dustin ML (2014) The immunological synapse. Cancer Immunol Res 2 (11):1023–1033. doi:10.1158/2326-6066.CIR-14-0161

2. Ehrlich LI, Ebert PJ, Krummel MF, Weiss A, Davis MM (2002) Dynamics of p56lck translocation to the T cell immunological synapse following agonist and antagonist stimulation. Immunity 17 (6):809–822. doi:10.1016/s1074-7613(02)00481-8

3. Das V, Nal B, Dujeancourt A, Thoulouze MI, Galli T, Roux P, Dautry-Varsat A, Alcover A (2004) Activation-induced polarized recycling targets T cell antigen receptors to the immunological synapse; involvement of SNARE complexes. Immunity 20 (5):577–588. doi:10.1016/s1074-7613(04)00106-2

4. Bonello G, Blanchard N, Montoya MC, Aguado E, Langlet C, He HT, Nunez-Cruz S, Malissen M, Sanchez-Madrid F, Olive D, Hivroz C, Collette Y (2004) Dynamic recruitment of the adaptor protein LAT: LAT exists in two distinct intracellular pools and controls its own recruitment. J Cell Sci 117 (Pt 7):1009–1016. doi:10.1242/jcs.00968

5. Soares H, Lasserre R, Alcover A (2013) Orchestrating cytoskeleton and intracellular vesicle traffic to build functional immunological synapses. Immunol Rev 256 (1):118–132. doi:10.1111/imr.12110

6. Finetti F, Cassioli C, Baldari CT (2017) Transcellular communication at the immunological synapse: a vesicular traffic-mediated mutual exchange. F1000Res 6:1880. doi:10.12688/f1000research.11944.1

7. Onnis A, Baldari CT (2019) Orchestration of Immunological Synapse Assembly by Vesicular Trafficking. Front Cell Dev Biol 7:110. doi:10.3389/fcell.2019.00110

8. Vardhana S, Choudhuri K, Varma R, Dustin ML (2010) Essential role of ubiquitin and TSG101 protein in formation and function of the central supramolecular activation cluster. Immunity 32 (4):531–540. doi:10.1016/j.immuni.2010.04.005

9. Compeer EB, Kraus F, Ecker M, Redpath G, Amiezer M, Rother N, Nicovich PR, Kapoor-Kaushik N, Deng Q, Samson GPB, Yang Z, Lou J, Carnell M, Vartoukian H, Gaus K, Rossy J (2018) A mobile endocytic network connects clathrin-independent receptor endocytosis to recycling and promotes T cell activation. Nat Commun 9 (1):1597. doi:10.1038/s41467-018-04088-w

10. Alcover A, Alarcon B, Di Bartolo V (2018) Cell Biology of T Cell Receptor Expression and Regulation. Annu Rev Immunol 36:103–125. doi:10.1146/annurev-immunol-042617-053429

11. Choudhuri K, Llodra J, Roth EW, Tsai J, Gordo S, Wucherpfennig KW, Kam LC, Stokes DL, Dustin ML (2014) Polarized release of T-cell-receptor-enriched microvesicles at the immunological synapse. Nature 507 (7490):118–123. doi:10.1038/nature12951

12. Pedersen LB, Rosenbaum JL (2008) Intraflagellar transport (IFT) role in ciliary assembly, resorption and signalling. Curr Top Dev Biol 85:23–61. doi:10.1016/S0070-2153(08)00802-8

13. Finetti F, Paccani SR, Riparbelli MG, Giacomello E, Perinetti G, Pazour GJ, Rosenbaum JL, Baldari CT (2009) Intraflagellar transport is required for polarized recycling of the TCR/CD3 complex to the immune synapse. Nat Cell Biol 11 (11):1332–1339. doi:10.1038/ncb1977

14. Finetti F, Patrussi L, Masi G, Onnis A, Galgano D, Lucherini OM, Pazour GJ, Baldari CT (2014) Specific recycling receptors are targeted to the immune synapse by the intraflagellar transport system. J Cell Sci 127 (Pt 9):1924–1937. doi:10.1242/jcs.139337

15. Martin-Cofreces NB, Baixauli F, Lopez MJ, Gil D, Monjas A, Alarcon B, Sanchez-Madrid F (2012) End-binding protein 1 controls signal propagation from the T cell receptor. EMBO J 31 (21):4140–4152. doi:10.1038/emboj.2012.242

16. Larghi P, Williamson DJ, Carpier JM, Dogniaux S, Chemin K, Bohineust A, Danglot L, Gaus K, Galli T, Hivroz C (2013) VAMP7 controls T cell activation by regulating the recruitment and phosphorylation of vesicular Lat at TCR-activation sites. Nat Immunol 14 (7):723–731. doi:10.1038/ni.2609

17. Onnis A, Finetti F, Patrussi L, Gottardo M, Cassioli C, Spano S, Baldari CT (2015) The small GTPase Rab29 is a common regulator of immune synapse assembly and ciliogenesis. Cell Death Differ 22 (10):1687–1699. doi:10.1038/cdd.2015.17

18. Finetti F, Patrussi L, Galgano D, Cassioli C, Perinetti G, Pazour GJ, Baldari CT (2015) The small GTPase Rab8 interacts with VAMP-3 to regulate the delivery of recycling T-cell receptors to the immune synapse. J Cell Sci 128 (14):2541–2552. doi:10.1242/jcs.171652

19. Bouchet J, Del Rio-Iniguez I, Lasserre R, Aguera-Gonzalez S, Cuche C, Danckaert A, McCaffrey MW, Di Bartolo V, Alcover A (2016) Rac1-Rab11-FIP3 regulatory hub coordinates vesicle traffic with actin remodeling and T-cell activation. EMBO J 35 (11):1160–1174. doi:10.15252/embj.201593274

20. Stephen LA, ElMaghloob Y, McIlwraith MJ, Yelland T, Castro Sanchez P, Roda-Navarro P, Ismail S (2018) The Ciliary Machinery Is Repurposed for T Cell Immune Synapse Trafficking of LCK. Dev Cell 47 (1):122–132 e124. doi:10.1016/j.devcel.2018.08.012

21. Zucchetti AE, Bataille L, Carpier JM, Dogniaux S, San Roman-Jouve M, Maurin M, Stuck MW, Rios RM, Baldari CT, Pazour GJ, Hivroz C (2019) Tethering of vesicles to the Golgi by GMAP210 controls LAT delivery to the immune synapse. Nat Commun 10 (1):2864. doi:10.1038/s41467-019-10891-w

22. de la Roche M, Ritter AT, Angus KL, Dinsmore C, Earnshaw CH, Reiter JF, Griffiths GM (2013) Hedgehog signaling controls T cell killing at the immunological synapse. Science 342 (6163):1247–1250. doi:10.1126/science.1244689

23. Gawden-Bone CM, Frazer GL, Richard AC, Ma CY, Strege K, Griffiths GM (2018) PIP5 Kinases Regulate Membrane Phosphoinositide and Actin Composition for Targeted Granule Secretion by Cytotoxic Lymphocytes. Immunity 49 (3):427–437 e424. doi:10.1016/j.immuni.2018.08.017

24. Baldari CT, Rosenbaum J (2010) Intraflagellar transport: it’s not just for cilia anymore. Curr Opin Cell Biol 22 (1):75–80. doi:10.1016/j.ceb.2009.10.010

25. de la Roche M, Asano Y, Griffiths GM (2016) Origins of the cytolytic synapse. Nat Rev Immunol 16 (7):421–432. doi:10.1038/nri.2016.54

26. Cassioli C, Baldari CT (2019) A Ciliary View of the Immunological Synapse. Cells 8 (8). doi:10.3390/cells8080789

27. Nachury MV (2018) The molecular machines that traffic signaling receptors into and out of cilia. Curr Opin Cell Biol 51:124–131. doi:10.1016/j.ceb.2018.03.004

28. Nakayama K, Katoh Y (2018) Ciliary protein trafficking mediated by IFT and BBSome complexes with the aid of kinesin-2 and dynein-2 motors. J Biochem 163 (3):155–164. doi:10.1093/jb/mvx087

29. Mourao A, Nager AR, Nachury MV, Lorentzen E (2014) Structural basis for membrane targeting of the BBSome by ARL6. Nat Struct Mol Biol 21 (12):1035–1041. doi:10.1038/nsmb.2920

30. Follit JA, Tuft RA, Fogarty KE, Pazour GJ (2006) The intraflagellar transport protein IFT20 is associated with the Golgi complex and is required for cilia assembly. Mol Biol Cell 17 (9):3781–3792. doi:10.1091/mbc.e06-02-0133

31. Baldari CT, Macchia G, Telford JL (1992) Interleukin-2 promoter activation in T-cells expressing activated Ha-ras. J Biol Chem 267 (7):4289–4291

32. Haeussler M, Schonig K, Eckert H, Eschstruth A, Mianne J, Renaud JB, Schneider-Maunoury S, Shkumatava A, Teboul L, Kent J, Joly JS, Concordet JP (2016) Evaluation of off-target and on-target scoring algorithms and integration into the guide RNA selection tool CRISPOR. Genome Biol 17 (1):148. doi:10.1186/s13059-016-1012-2

33. Ran FA, Hsu PD, Wright J, Agarwala V, Scott DA, Zhang F (2013) Genome engineering using the CRISPR-Cas9 system. Nat Protoc 8 (11):2281–2308. doi:10.1038/nprot.2013.143

34. Chicaybam L, Sodre AL, Curzio BA, Bonamino MH (2013) An efficient low cost method for gene transfer to T lymphocytes. PLoS One 8 (3):e60298. doi:10.1371/journal.pone.0060298

35. Manders EM, Stap J, Brakenhoff GJ, van Driel R, Aten JA (1992) Dynamics of three-dimensional replication patterns during the S-phase, analysed by double labelling of DNA and confocal microscopy. J Cell Sci 103 (Pt 3):857–862

36. Esquerre M, Tauzin B, Guiraud M, Muller S, Saoudi A, Valitutti S (2008) Human regulatory T cells inhibit polarization of T helper cells toward antigen-presenting cells via a TGF-beta-dependent mechanism. Proc Natl Acad Sci USA 105 (7):2550–2555. doi:10.1073/pnas.0708350105

37. Obino D, Farina F, Malbec O, Saez PJ, Maurin M, Gaillard J, Dingli F, Loew D, Gautreau A, Yuseff MI, Blanchoin L, Thery M, Lennon-Dumenil AM (2016) Actin nucleation at the centrosome controls lymphocyte polarity. Nat Commun 7:10969. doi:10.1038/ncomms10969

38. Prasai A, Schmidt Cernohorska M, Ruppova K, Niederlova V, Andelova M, Draber P, Stepanek O, Huranova M (2020) The BBSome assembly is spatially controlled by BBS1 and BBS4 in human cells. J Biol Chem 295 (42):14279–14290. doi:10.1074/jbc.RA120.013905

39. Wingfield JL, Lechtreck KF, Lorentzen E (2018) Trafficking of ciliary membrane proteins by the intraflagellar transport/BBSome machinery. Essays Biochem 62 (6):753–763. doi:10.1042/EBC20180030

40. Ansley SJ, Badano JL, Blacque OE, Hill J, Hoskins BE, Leitch CC, Kim JC, Ross AJ, Eichers ER, Teslovich TM, Mah AK, Johnsen RC, Cavender JC, Lewis RA, Leroux MR, Beales PL, Katsanis N (2003) Basal body dysfunction is a likely cause of pleiotropic Bardet-Biedl syndrome. Nature 425 (6958):628–633. doi:10.1038/nature02030

41. Katsanis N (2004) The oligogenic properties of Bardet-Biedl syndrome. Hum Mol Genet 13 Spec No 1:R65–71. doi:10.1093/hmg/ddh092

42. Tobin JL, Beales PL (2009) The nonmotile ciliopathies. Genet Med 11 (6):386–402. doi:10.1097/GIM.0b013e3181a02882

43. Burakov AV, Nadezhdina ES (2013) Association of nucleus and centrosome: magnet or velcro? Cell Biol Int 37 (2):95–104. doi:10.1002/cbin.10016

44. Burke B (2019) Chain reaction: LINC complexes and nuclear positioning. F1000Res 8. doi:10.12688/f1000research.16877.1

45. Rajgor D, Shanahan CM (2013) Nesprins: from the nuclear envelope and beyond. Expert Rev Mol Med 15:e5. doi:10.1017/erm.2013.6

46. Bello-Gamboa A, Velasco M, Moreno S, Herranz G, Ilie R, Huetos S, Davila S, Sancheza A, De La Serna JB, Calvo V, Izquierdo M (2020) Actin reorganization at the centrosomal area and the immune synapse regulates polarized secretory traffic of multivesicular bodies in T lymphocytes. J Extracell Vesicles 9 (1). doi:10.1080/20013078.2020.1759926

47. Seaman MN, Gautreau A, Billadeau DD (2013) Retromer-mediated endosomal protein sorting: all WASHed up! Trends Cell Biol 23 (11):522–528. doi:10.1016/j.tcb.2013.04.010

48. Gomez TS, Billadeau DD (2009) A FAM21-containing WASH complex regulates retromer-dependent sorting. Dev Cell 17 (5):699–711. doi:10.1016/j.devcel.2009.09.009

49. Farina F, Gaillard J, Guerin C, Coute Y, Sillibourne J, Blanchoin L, Thery M (2016) The centrosome is an actin-organizing centre. Nat Cell Biol 18 (1):65–75. doi:10.1038/ncb3285

50. Vora SM, Phillips BT (2016) The benefits of local depletion: The centrosome as a scaffold for ubiquitin-proteasome-mediated degradation. Cell Cycle 15 (16):2124–2134. doi:10.1080/15384101.2016.1196306

51. Ibanez-Vega J, Del Valle Batalla F, Saez JJ, Soza A, Yuseff MI (2019) Proteasome Dependent Actin Remodeling Facilitates Antigen Extraction at the Immune Synapse of B Cells. Front Immunol 10:225. doi:10.3389/fimmu.2019.00225

52. Bard JAM, Goodall EA, Greene ER, Jonsson E, Dong KC, Martin A (2018) Structure and Function of the 26S Proteasome. Annu Rev Biochem 87:697–724. doi:10.1146/annurev-biochem-062917-011931

53. Hsu MT, Guo CL, Liou AY, Chang TY, Ng MC, Florea BI, Overkleeft HS, Wu YL, Liao JC, Cheng PL (2015) Stage-Dependent Axon Transport of Proteasomes Contributes to Axon Development. Dev Cell 35 (4):418–431. doi:10.1016/j.devcel.2015.10.018

54. Liu YP, Tsai IC, Morleo M, Oh EC, Leitch CC, Massa F, Lee BH, Parker DS, Finley D, Zaghlou NA, Franco B, Katsanis N (2014) Ciliopathy proteins regulate paracrine signaling by modulating proteasomal degradation of mediators. J Clin Invest 124 (5):2059–2070. doi:10.1172/Jci71898

55. Starks RD, Beyer AM, Guo DF, Boland L, Zhang QH, Sheffield VC, Rahmouni K (2015) Regulation of Insulin Receptor Trafficking by Bardet Biedl Syndrome Proteins. Plos Genetics 11 (6). doi:10.1371/journal.pgen.1005311

56. Guo DF, Cui HX, Zhang QH, Morgan DA, Thedens DR, Nishimura D, Grobe JL, Sheffield VC, Rahmouni K (2016) The BBSome Controls Energy Homeostasis by Mediating the Transport of the Leptin Receptor to the Plasma Membrane. Plos Genetics 12 (2). doi:10.1371/journal.pgen.1005890

57. Jin H, White SR, Shida T, Schulz S, Aguiar M, Gygi SP, Bazan JF, Nachury MV (2010) The conserved Bardet-Biedl syndrome proteins assemble a coat that traffics membrane proteins to cilia. Cell 141 (7):1208–1219. doi:10.1016/j.cell.2010.05.015

58. Nozaki S, Katoh Y, Kobayashi T, Nakayama K (2018) BBS1 is involved in retrograde trafficking of ciliary GPCRs in the context of the BBSome complex. PLoS One 13 (3):e0195005. doi:10.1371/journal.pone.0195005

59. Ye F, Nager AR, Nachury MV (2018) BBSome trains remove activated GPCRs from cilia by enabling passage through the transition zone. J Cell Biol 217 (5):1847–1868. doi:10.1083/jcb.201709041

60. Stinchcombe JC, Griffiths GM (2014) Communication, the centrosome and the immunological synapse. Philos Trans R Soc Lond B Biol Sci 369 (1650). doi:10.1098/rstb.2013.0463

61. Bustos-Moran E, Blas-Rus N, Martin-Cofreces NB, Sanchez-Madrid F (2016) Orchestrating Lymphocyte Polarity in Cognate Immune Cell-Cell Interactions. Int Rev Cell Mol Biol 327:195–261. doi:10.1016/bs.ircmb.2016.06.004

62. Xu Q, Zhang Y, Wei Q, Huang Y, Li Y, Ling K, Hu J (2015) BBS4 and BBS5 show functional redundancy in the BBSome to regulate the degradative sorting of ciliary sensory receptors. Sci Rep 5:11855. doi:10.1038/srep11855

63. Langousis G, Shimogawa MM, Saada EA, Vashisht AA, Spreafico R, Nager AR, Barshop WD, Nachury MV, Wohlschlegel JA, Hill KL (2016) Loss of the BBSome perturbs endocytic trafficking and disrupts virulence of Trypanosoma brucei. Proc Natl Acad Sci USA 113 (3):632–637. doi:10.1073/pnas.1518079113

64. Desai PB, Stuck MW, Lv B, Pazour GJ (2020) Ubiquitin links smoothened to intraflagellar transport to regulate Hedgehog signaling. J Cell Biol 219 (7). doi:10.1083/jcb.201912104

65. Hao YH, Doyle JM, Ramanathan S, Gomez TS, Jia D, Xu M, Chen ZJ, Billadeau DD, Rosen MK, Potts PR (2013) Regulation of WASH-dependent actin polymerization and protein trafficking by ubiquitination. Cell 152 (5):1051–1064. doi:10.1016/j.cell.2013.01.051

66. Watanabe Y, Sasahara Y, Ramesh N, Massaad MJ, Yeng Looi C, Kumaki S, Kure S, Geha RS, Tsuchiya S (2013) T-cell receptor ligation causes Wiskott-Aldrich syndrome protein degradation and F-actin assembly downregulation. J Allergy Clin Immunol 132 (3):648–655 e641. doi:10.1016/j.jaci.2013.03.046

67. Joseph N, Biber G, Fried S, Reicher B, Levy O, Sabag B, Noy E, Barda-Saad M (2017) A conformational change within the WAVE2 complex regulates its degradation following cellular activation. Sci Rep 7:44863. doi:10.1038/srep44863

68. Didier C, Merdes A, Gairin JE, Jabrane-Ferrat N (2008) Inhibition of proteasome activity impairs centrosome-dependent microtubule nucleation and organization. Mol Biol Cell 19 (3):1220–1229. doi:10.1091/mbc.e06-12-1140

69. Forsythe E, Beales PL (2013) Bardet-Biedl syndrome. Eur J Hum Genet 21 (1):8–13. doi:10.1038/ejhg.2012.115

70. Tsang SH, Aycinena ARP, Sharma T (2018) Ciliopathy: Bardet-Biedl Syndrome. Adv Exp Med Biol 1085:171–174. doi:10.1007/978-3-319-95046-4_33

71. Mourao A, Christensen ST, Lorentzen E (2016) The intraflagellar transport machinery in ciliary signaling (vol 41, pg 98, 2016). Curr Opin Struct Biol 41:255–255. doi: 10.1016/j.sbi.2016.07.018

72. Tsyklauri O, Niederlova V, Forsythe E, Drobek A, Prasai A, Sparks K, Trachtulek Z, Beales P, Huranova M, Stepanek O. Altered hematopoietic system and self-tolerance in Bardet-Biedl Syndrome. bioRxiv 962886; doi: https://doi.org/10.1101/2020.02.24.962886.

73. Mestas J, Hughes CCW (2004) Of mice and not men: Differences between mouse and human immunology. J Immunol 172 (5):2731–2738. doi:DOI 10.4049/jimmunol.172.5.2731

74. Cu Bjornson-Hooper ZB, Fragiadakis GK, Spitzer MH, Madhireddy D, McIlwain D, Nolan GP. A comprehensive atlas of immunological differences between humans, mice and non-human primates. bioRxiv 574160; doi: https://doi.org/10.1101/574160

