## Supplemental figures and tables for "The Bardet-Biedl Syndrome complex component BBS1 regulates proteasome-dependent F-actin clearance from the centrosome to enable its translocation to the T cell immune synapse"

**SUPPLEMENTAL FIGURE LEGENDS**

**Fig. S1** (**a**) Quantitative RT-PCR of the BBSome subunits BBS-1, -2, -4, -5, -7, -8, -9 and -18 in Jurkat cells, primary T cells and BJ-5at cells (n≥3). (**b**) Immunoblot analysis of BBS1 in lysates of Jurkat cells transduced with lentiviral particles containing a non-targeting control shRNA (ctr) or a shRNA specific for BBS1 (J KD) (*left*), or in lysates of control (ctr) primary and BBS1 KO (T KO) primary T cells gene-edited by CRISPR-Cas9 technology (*middle*) or in lysates of control (ctr) and BBS1 KO (J KO) Jurkat cells gene-edited by CRISPR-Cas9 technology (*right*) (n≥3). Actin was used as loading control. Representative immunoblots are shown. The migration of molecular mass markers is shown for each filter. The quantification of the relative protein expression normalized to actin (mean±SD, n≥3, paired two-tailed Student's *t*-test). (**c**) Flow cytometric analysis of CD3ε in Jurkat cells transduced with lentiviral particles containing a non-targeting control shRNA (ctr) or a shRNA specific for BBS1 (J KD) (*left*), in control (ctr) and BBS1 KO (T KO) primary T cells (*middle*), or control (ctr) and BBS1 KO (J KO) Jurkat cells (*right*) (n≥3). The mean fluorescence intensity (MFI) is reported in each representative FACS profile. (**d**) Quantitative RT-PCR of the BBSome subunits BBS-1, -2, -4, -5, -7, -8, -9 and -18 in Jurkat cells transduced with lentiviral particles containing a non-targeting control shRNA (ctr) or a shRNA specific for BBS1 (J KD). (**e**) Immunoblot analysis of BBS1 in lysates of control (ctr) or BBS1 KO (J KO) Jurkat cells, the latter transfected with either empty vector (ctr, J KO) or the same vector encoding wild-type BBS1 (J KO+BBS1) (n=3). The migration of molecular mass markers is shown for each filter. The quantification of the relative protein expression normalized to actin is shown on the right. (**f**) Flow cytometric analysis of GFP in control (ctr) Jurkat cells either untransfected (neg ctr) or transfected with empty vector (ctr+GFP), or BBS1 KO (J KO) Jurkat cells transfected with either empty vector (J KO+GFP) or the same vector encoding wild-type BBS1 (J KO+BBS1-GFP) (n=3). The percentages (%) of GFP$^{+}$/propidium iodide$^{-}$ cells are shown in each representative dot plot. The data are expressed as mean±SD. ***P≤0.001; **P≤0.01; *P≤0.05.

**Fig. S2** (**a**) Immunofluorescence analysis of BBS1-GFP expressing Jurkat cells and stained for CEP290 (pericentrosomal marker), Rab11 (recycling endosomes) and Rab7 (late endosomes). The histograms show the intensity profiles along the white lines within the selected area in the overlay images for each channel. The raw pixel intensity signals were normalized to maximum intensity pixel of each channel (% max grey value). A quantification using Mander's coefficient of the co-localization of BBS1-GFP with these markers is shown in Figure 1C. At least 20 Jurkat cells were analyzed for each marker (n≥3). Representative images (medial optical sections) are shown. (**b**) Immunofluorescence analysis of PCNT and CD3ζ localization in conjugates of control (ctr) and BBS1KD (J KD) Jurkat cells with Raji B cells (APC), in the absence or presence of SEE (n=3). Size bar, 5 µm.

**Fig. S3** (**a,b**) Immunofluorescence analysis of CD3ζ, PTyr and γ–tubulin in conjugates of control (ctr) and BBS1KD (J KD) Jurkat cells in the absence of SEE (**a**) or in conjugates of control (ctr) and BBS1KO (T KO) primary T cells with Raji B cells (APC) in the absence of SAg (**b**) (n≥3). Size bar, 5 µm. (**c**) Time course flow cytometric analysis of protein tyrosine phosphorylation in conjugates of control (ctr) and BBS1 KO (J KO) Jurkat cells with SEE-loaded Raji B cells (APC). Raji cells were labelled with DiO and the analysis was carried out gating on DiO-negative cells. The histogram shows the mean fluorescence intensity (MFI) of PTyr$^{+}$/DiO$^{-}$ cells at different time points (n=3, Mann-Whitney test). **P≤0.01; *P≤0.05.

**Fig. S4** (**a**) Representative image of Jurkat cells conjugated with Raji B cells for 15 min and co-stained with anti-nesprin-2 and anti-PCNT antibodies. Jurkat T cell was magnified to depict the parameters used for quantification. A line from the center of the centrosome (PCNT) forming an angle of 90° with the tangent of the nuclear membrane (nesprin-2) was drawn to measure the distance between nucleus and centrosome (μm) in Jurkat and primary T cells as shown in Fig.3a,b and Fig.5b,f. In cells showing a weak nesprin-2 staining the nucleus-centrosome distance was further confirmed by overlaying the PCNT (or PCM1) fluorescence to the DIC image, in which the nuclear contour is easily identified. (**b**) Representative image of Jurkat cells conjugated with Raji B cells (APC) for 15 min and co-stained for F-actin and PCNT antibodies. The representative Jurkat T cell has been magnified to depict the parameters used for quantification. A 4.5-μm diameter circle centered around the centrosome of Jurkat cells (or 2.85 μm for primary T cells) indicates the centrosomal area used for the colocalization analyses shown in Fig.4a-d, Fig.5i-j and Fig.6a,b. Size bar, 5 μm.

**Fig. S5** (**a**) Immunofluorescence analysis of conjugates of control (ctr) and BBS1KD (J KD) Jurkat cells with SEE-loaded Raji B cells (APC) co-stained with anti-PCM-1 and anti-γ–tubulin antibodies. Representative images (medial optical sections) are shown. Size bar, 5 μm. (**b,d-f**) Quantification using Mander's coefficient of the weighted colocalization of PCM-1 with γ–tubulin (**b**), centrosomal F-actin (**d**), 19S RP (**e**) and Ubiquitin (**f**) is shown. The parameters used for this quantification as schematized in Fig.S4B. The histograms show the quantification in conjugates of control (ctr) and BBS1KD (J KD) Jurkat cells with Raji B cells (APC) in the absence or presence of SEE (≥ 10 cells/sample, n=3, Kruskal-Wallis test). (**c**) Measurement of the distance (μm) of PCM1 from the nucleus in conjugates of control (ctr) and BBS1KD (J KD) Jurkat cells with Raji B cells (APC) in the absence or presence of SEE (≥ 10 cells/sample, n=3, Kruskal-Wallis test). The data are expressed as mean±SD. **** P ≤ 0.0001; **P≤0.01; *P≤0.05.

**Fig. S6** (**a,b**) Immunofluorescence analysis of centrosomal F-actin$^+$ in 15 min conjugates of control (ctr) and BBS1KD (J KD) Jurkat cells with Raji B cells (APC) in the absence or presence of SEE, co-stained for Rab5 (**a**) and Rab11 (**b**). The parameters used for this quantification as schematized in Fig.S4B. The histograms show the co-localization of F-actin on individual dots (*left*) and the quantification of F-actin$^+$ dots positive for Rab5 or Rab11 (*right*) were shown below each panel. Measurements were taken on 10 conjugates from 1 experiment and a mean of 15 dots per cell were analyzed. Size bar, 5 µm. The data are expressed as mean±SD.

**Fig. S7** (**a**) Immunoblot analysis of Ubiquitin in lysates of control (ctr) Jurkat cells either untreated (DMSO) or pre-treated with the proteasome inhibitors MG132 and epoxomicin (n=3). Actin was used as loading control. (**b**) Viability (%) of control (ctr) Jurkat cells either untreated (DMSO) or pre-treated with the proteasome inhibitors MG132 and epoxomicin measured using trypan blue exclusion (n=3). The data are expressed as mean±SD. (**c**) Immunoblot analysis of ubiquitin in lysates of control (ctr) and BBS1KD (J KD) Jurkat cells. Actin was used as loading control (n≥3). The migration of molecular mass markers is indicated.

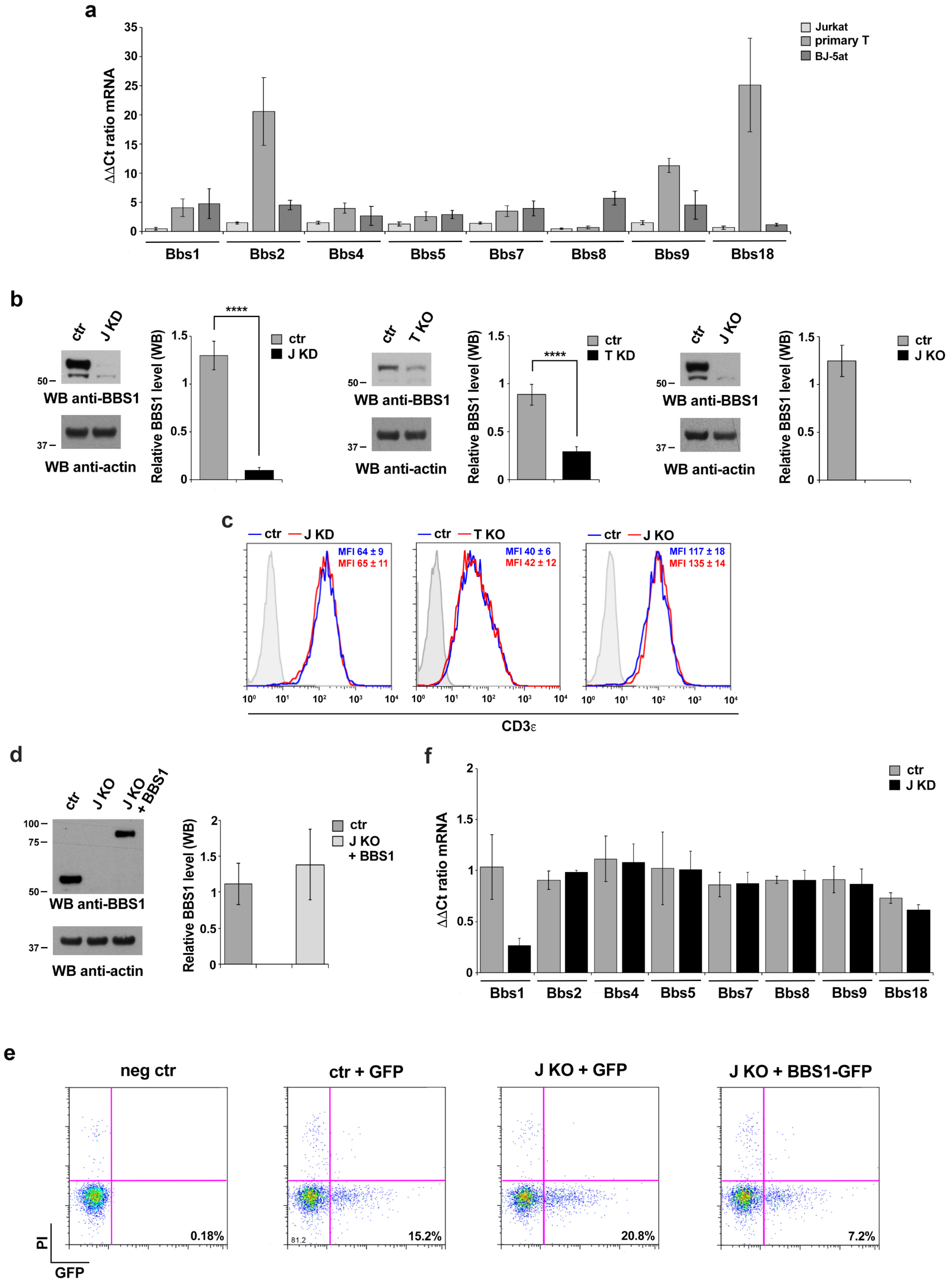

a
ΔΔCt ratio mRNA
Jurkat
primary T
BJ-5at
Bbs1
Bbs2
Bbs4
Bbs5
Bbs7
Bbs8
Bbs9
Bbs18
b
ctr
J KD
WB anti-BBS1
WB anti-actin
Relative BBS1 level (WB)
****
T KO
J KO
c
MFI 64 ± 9
MFI 65 ± 11
MFI 40 ± 6
MFI 42 ± 12
MFI 117 ± 18
MFI 135 ± 14
CD3ε
d
J KO + BBS1
f
e
neg ctr
ctr + GFP
J KO + GFP
J KO + BBS1-GFP
0.18%
15.2%
20.8%
7.2%
PI
GFP


FIGURE S1

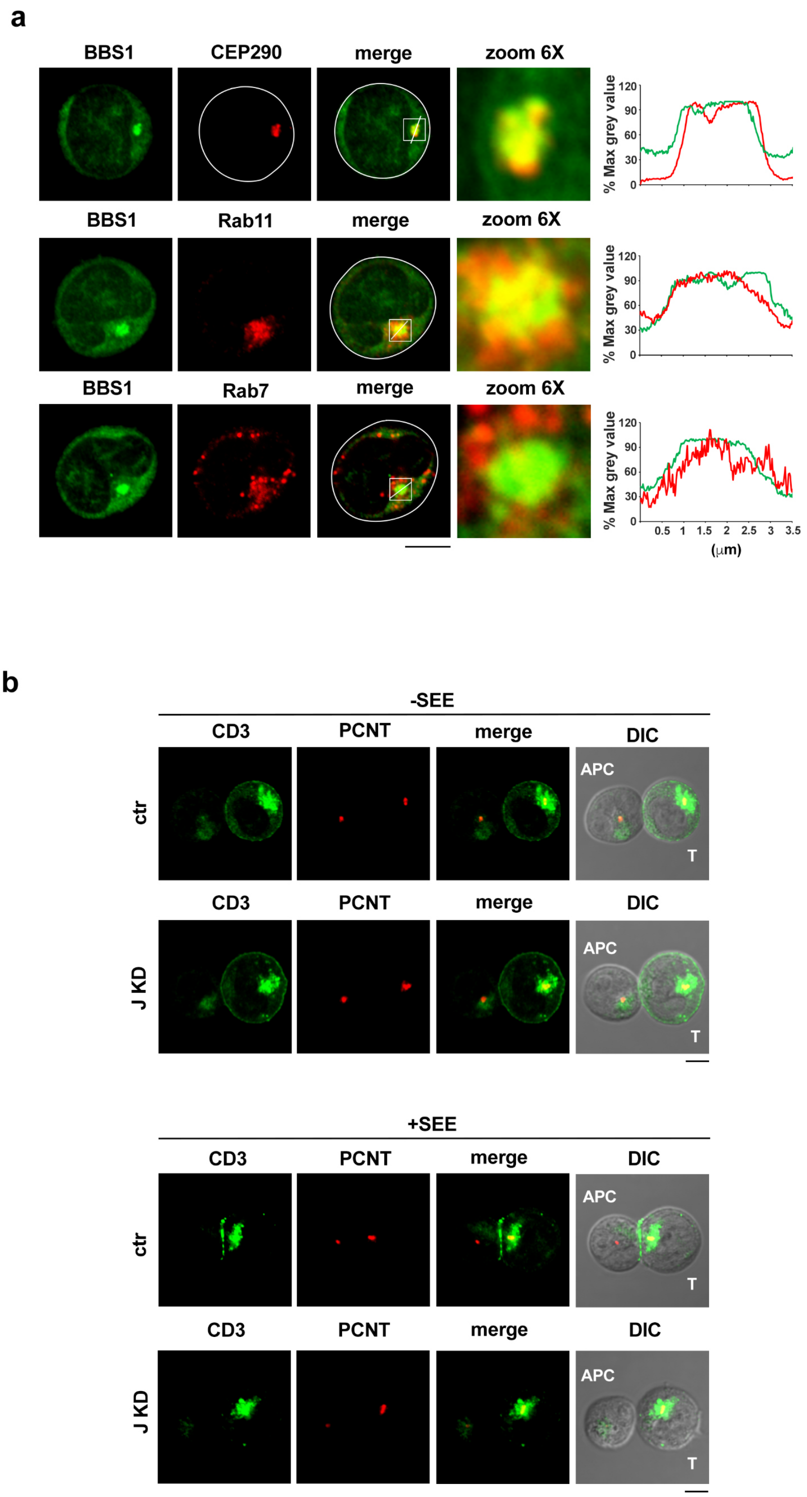

a
BBS1
CEP290
merge
zoom 6X
% Max grey value
Rab11
Rab7
0.5 1 1.5 2 2.5 3 3.5
(µm)
b
-SEE
+SEE
CD3
PCNT
DIC
APC
T
ctr
J KD


### FIGURE S2

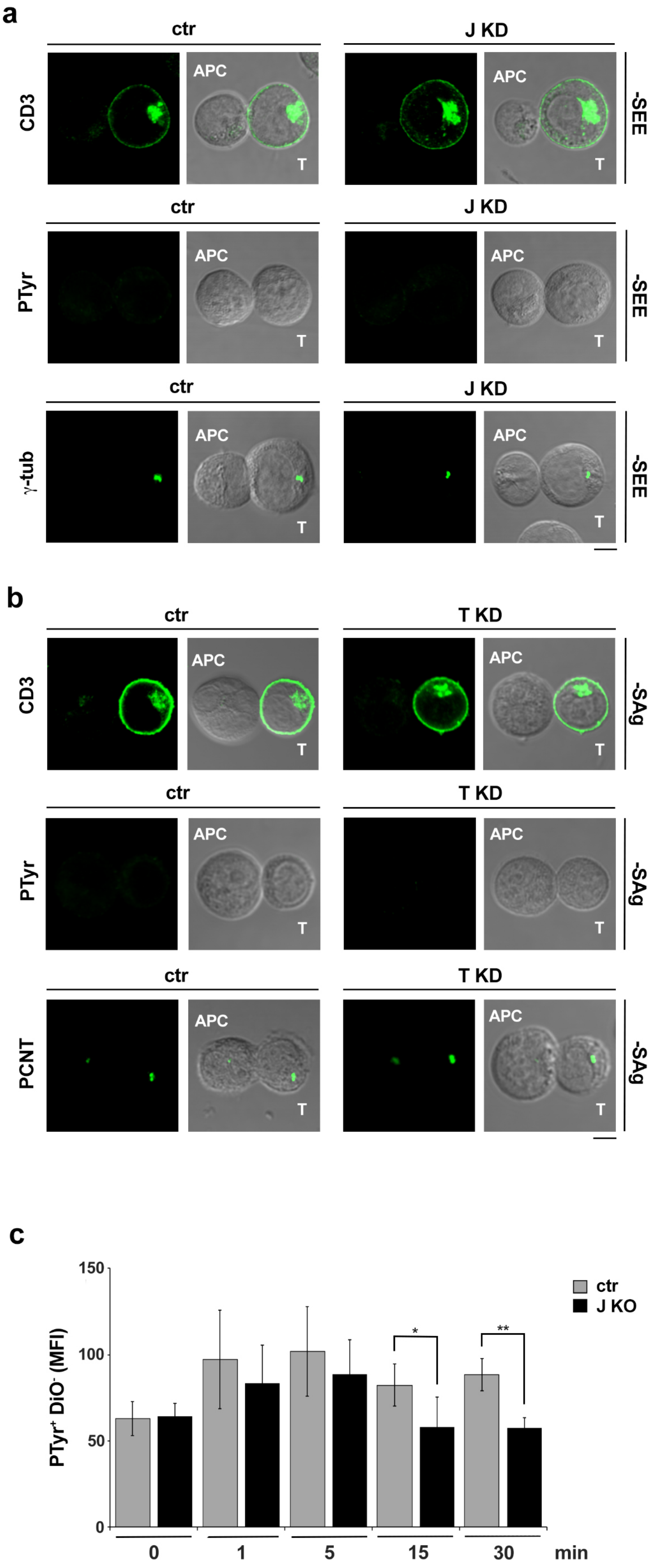
a
ctr
J KD
CD3
PTyr
γ-tub
APC
T
-SEE
b
T KD
PCNT
-SAg
c
150
100
50
0
PTyr$^+$ DiO$^-$ (MFI)
ctr
J KO
*
**
1
5
15
30
min

### FIGURE S3

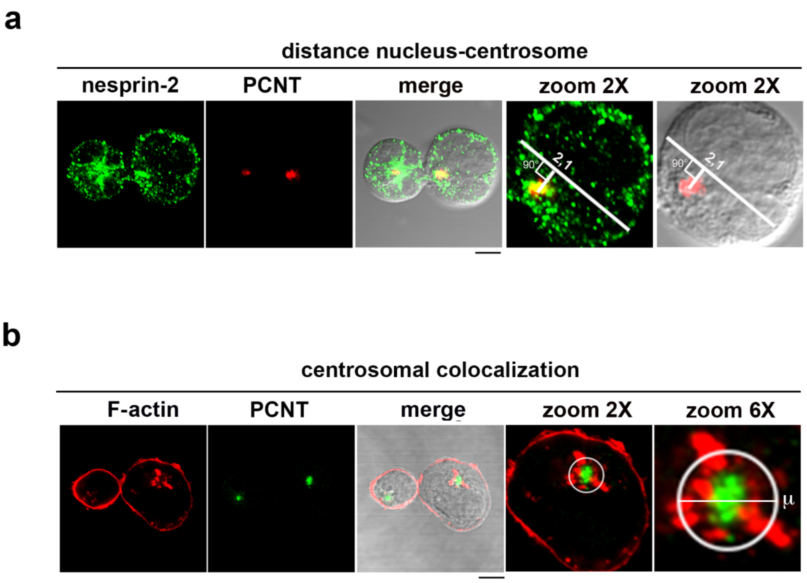
a
distance nucleus-centrosome
nesprin-2
PCNT
merge
zoom 2X
zoom 2X
90°
2,1
b
centrosomal colocalization
F-actin
PCNT
merge
zoom 2X
zoom 6X
μ

### FIGURE S4

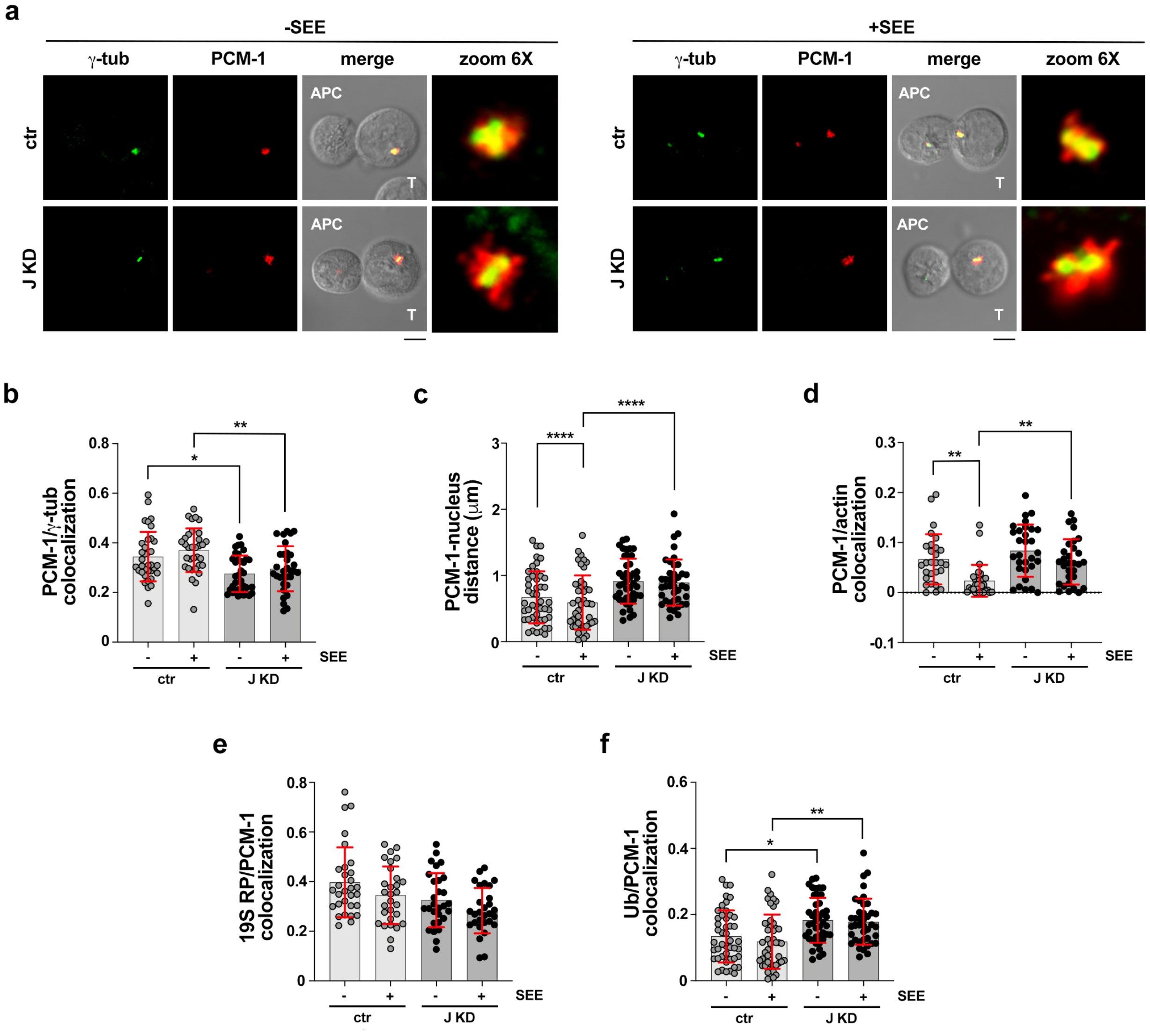

a
-SEE
+SEE
γ-tub
PCM-1
merge
zoom 6X
ctr
J KD
APC
T
b
PCM-1/γ-tub colocalization
c
PCM-1-nucleus distance (μm)
d
PCM-1/actin colocalization
e
19S RP/PCM-1 colocalization
f
Ub/PCM-1 colocalization
SEE
ctr
J KD


FIGURE S5

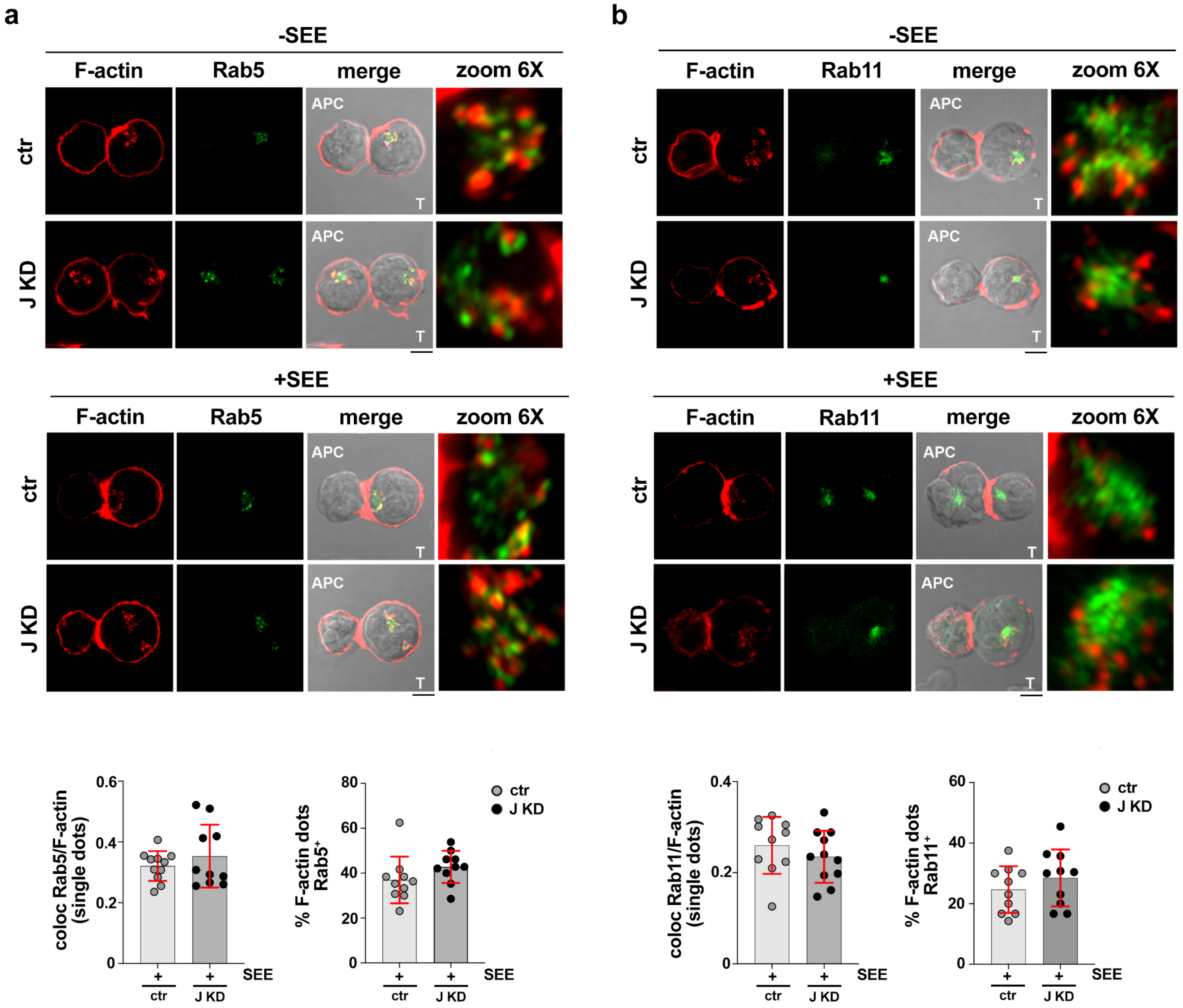
a
-SEE
F-actin
Rab5
merge
zoom 6X
ctr
J KD
APC
T
+SEE
coloc Rab5/F-actin (single dots)
% F-actin dots Rab5+
ctr
J KD
SEE
b
Rab11
coloc Rab11/F-actin (single dots)
% F-actin dots Rab11+

### FIGURE S6

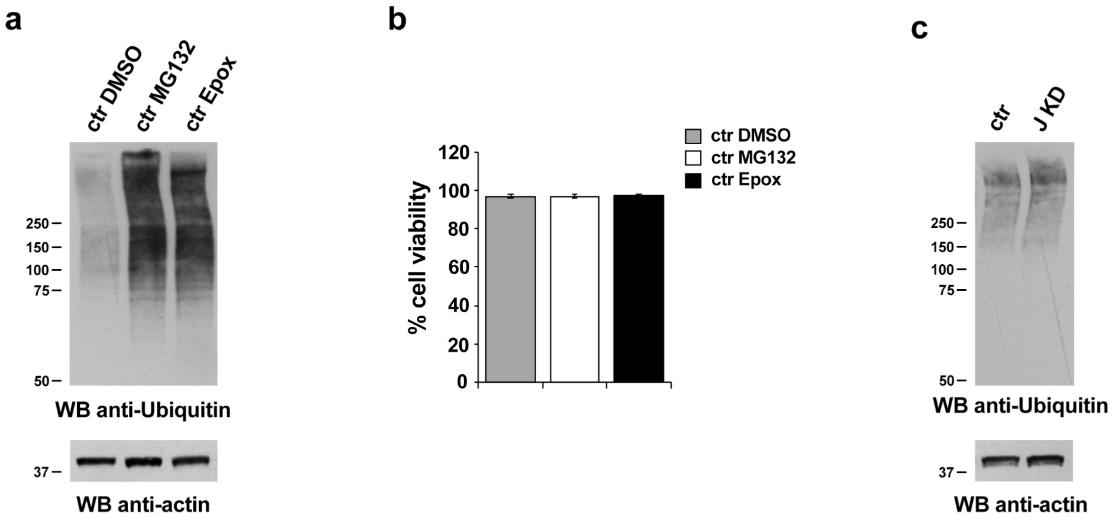
a
ctr DMSO
ctr MG132
ctr Epox
250
150
100
75
50
WB anti-Ubiquitin
37
WB anti-actin
b
% cell viability
120
100
80
60
40
20
0
ctr DMSO
ctr MG132
ctr Epox
c
ctr
J KD
250
150
100
75
50
WB anti-Ubiquitin
37
WB anti-actin

**FIGURE S7**

#### SUPPLEMENTAL TABLES

##### Table S1. List of the primers used in this study

| Oligo name | Sequence | Description |
|---|---|---|
| common reverse primer | AGCACCGACTCGGTGCCACT | sgRNA production |
| GFP sgRNA | ttaatacgactcactataggGGGCGAGGAG CTGTTCACCGgttttagagctagaaatagc | sgRNA production |
| hBBS1 gRNA1 | ttaatacgactcactataggGGGCTTTCGG TCATCACCAGgttttagagctagaaatagc | sgRNA production |
| hBBS1 gRNA2 | ttaatacgactcactataggGGATGCGCAC TACGACCCAAgttttagagctagaaatagc | sgRNA production |
| Bbs1 fw | AATTCGAAGTGGTTGGATGC | primer |
| Bbs1 rv | ATGTTGCTCCATGAGGAAGG | primer |
| Bbs2 fw | TTCCCCTCTTGCGATTATTG | primer |
| Bbs2 rv | GTCACACAAGGCCAAGGAAT | primer |
| Bbs4 fw | ACCACTTCAACCAGCAAACC | primer |
| Bbs4 rv | GGCTTTGTGAACTGGGATGT | primer |
| Bbs5 fw | CGGATGCTTTTGTGGCTTAT | primer |
| Bbs5 rv | CCAAAGTCCCTGTAGGGTGA | primer |
| Bbs7 fw | CCAGTACGCAAGTGGGAAAT | primer |
| Bbs7 rv | CTTCCACCATTCCGTCATCT | primer |
| Bbs8 fw | AGAGGCAGCTGATGTCTGGT | primer |
| Bbs8 rv | GCGTGGTTGTTGTTGTTGAC | primer |
| Bbs9 fw | CCCCACATTCCTGTAGCAGT | primer |
| Bbs9 rv | AGAAGGATCTGTCCCCAGGT | primer |
| Bbs18 fw | ACCATCTCGACTCACTGCAA | primer |
| Bbs18 rv | TGAGATTTAAGGGCTGGGCA | primer |
| HPRT1 fw | AGATGGTCAAGGTCGCAAG | primer |
| HPRT1 rv | GTATTCATTATAGTCAAGGGCATATC | primer |

**Table S2. List of the antibodies used in this study**

| Antibody | Host Species | Cat.No. | Source | Dilution WB | Dilution IF | Dilution FC |
|---|---|---|---|---|---|---|
| AF555 phalloidin | | A34055 | Invitrogen | - | 1:100<br>1:50 | - |
| anti-actin | mouse | MAB1501 | EMD Millipore | 1:10000 | - | - |
| anti-BBS1 | rabbit | ab166613 | AbCam | 1:1000 | - | - |
| anti-CD3-ε | mouse | 317308 | Biolegend | - | - | 1:100 |
| anti-CD3-ζ | mouse | sc-1239 | Santa Cruz | - | 1:100 | - |
| anti-CEP131 | rabbit | ab99379 | AbCam | - | 1:300 | - |
| anti-CEP290 | rabbit | ab84870 | AbCam | - | 1:300 | - |
| anti-dynein | mouse | MAB1618 | Merck Millipore | 1:500 | - | - |
| anti-γ-tubulin | mouse | T6557 | Sigma Aldrich | 1:5000 | 1:200 | - |
| anti-GM130 | mouse | 610822 | BD | - | 1:100 | - |
| anti-GFP | mouse | A11120 | Invitrogen | - | 1:200 | - |
| anti-GFP | rabbit | A11122 | Invitrogen | 1:1000 | 1:200 | - |
| anti-nesprin 2 | mouse | NBP2-59944 | Novus | - | 1:50 | - |
| anti-P-Tyr | mouse | 05-1050 | Merck Millipore | - | 1:100 | 1:400 |
| anti-PCM-1 | mouse | sc-398365 | Santa Cruz | - | 1:400 | - |
| anti-PCM-1 | rabbit | A301-150A | Bethyl | - | 1:500 | - |
| anti-pericentrin | rabbit | ab4448 | AbCam | - | 1:200 | - |
| anti-Rab7 | mouse | sc-376362 | Santa Cruz | - | 1:50 | - |
| anti-Rab11a | rabbit | 2413 | Cell Signaling | - | 1:50 | - |
| anti-tubulin | mouse | 15115 | Cell Signaling | 1:2000 | - | - |
| anti-Ubiquitin | mouse | 3936 | Cell Signaling | 1:500 | - | - |
| anti-Ubiquitin | mouse | 04-263 | Merck Millipore | - | 1:50 | - |
| anti-WASH | rabbit | PA5-51731 | Invitrogen | 1:500 | 1:100 | - |
| anti-19S RP | rabbit | ab140450 | AbCam | 1:2000 | - | - |
| anti-19S RP | rabbit | ab3317 | AbCam | - | 1:100 | - |
